# Optogenetic inhibition of Gα signalling alters and regulates circuit functionality and early circuit formation

**DOI:** 10.1101/2023.05.06.539674

**Authors:** Jayde Lockyer, Andrew Reading, Silvia Vicenzi, Caroline Delandre, Owen Marshall, Robert Gasperini, Lisa Foa, John Y. Lin

**Affiliations:** Tasmanian School of Medicine, University of Tasmania, Tasmania, Australia; Current affiliation, Moores Cancer Center, School of Medicine, Division of Regenerative Medicine, University of California, San Diego, California, USA; Menzies Institute of Medical Research, University of Tasmania, Tasmania, Australia; School of Psychological Sciences, University of Tasmania, Tasmania, Australia

**Author notes:** Contact: John Y. Lin Tasmanian School of Medicine College of Health and Medicine Private Bag 34 University of Tasmania Medical Sciences 2 Building, 17 Liverpool Street, Hobart, TAS 7000 Australia.

**Keywords:** Optogenetics, G-protein signaling, axon guidance, courtship learning

## Abstract

Optogenetic techniques provide genetically targeted, spatially and temporally precise approaches to correlate cellular activities and physiological outcomes. In the nervous system, G-protein-coupled receptors (GPCRs) have essential neuromodulatory functions through binding extracellular ligands to induce intracellular signaling cascades. In this work, we develop and validate a new optogenetic tool that disrupt Gα_q_ signaling through membrane recruitment of a minimal Regulator of G-protein signaling (RGS) domain. This approach, **P**hoto-**i**nduced **M**odulation of **G**α protein – **I**nhibition of Gα_q_ (PiGM-Iq), exhibited potent and selective inhibition of Gα_q_ signaling. We alter the behavior of *C. elegans* and *Drosophila* with outcomes consistent with GPCR-Gα_q_ disruption. PiGM-Iq also changes axon guidance in culture dorsal root ganglia neurons in response to serotonin. PiGM-Iq activation leads to developmental deficits in zebrafish embryos and larvae resulting in altered neuronal wiring and behavior. By altering the choice of minimal RGS domain, we also show that this approach is amenable to Gα_i_ signaling.

## Introduction

G protein-coupled receptors (GPCRs) in the nervous system mediate the intracellular signaling of extracellular chemical and peptidergic ligands. In the canonical GPCR signaling pathway, binding of the ligand and the activation of the receptor leads to the GDP-GTP exchange on the Gα subunit and the dissociation of GTP-bound Gα and Gßγ dimeric complex. The Gα and Gßγ mediate further signal transduction within the cells. GPCRs associate the canonical Gα proteins, which are named Gα_q/11_, Gα_s_, Gα_i/o_, and G_12/13._ Each of these G proteins activates distinct signaling cascades for unique outcomes. Gα_q/11_ activates PLCß, Gα_s_ activates adenylate cyclase, Gα_i/o_ inhibits the activities of adenylate cyclase, and G_12/13_ activates guanine nucleotide exchange factors. Gαq_/11_ activation of PLCß leads to the catalytic breakdown of PIP_2_ into IP_3_ and DAG, which leads to the release of calcium from an intracellular store and the activation of protein kinase C. In the central nervous system, Gα_q/11_ activation is associated with mGluR-induced long-term depression ^1^ and neuropeptides, such as oxytocin and vasopressin, control social, innate, and emotive behavior ^2^. Gα_i/o_ is structurally related to Gα_q_ and binds to adenylate cyclase to mediate its inhibitory effect ^3^. The termination of the GPCR signaling is often associated with the binding of ß-arrestin and endocytosis of the receptor. Alternatively, proteins containing regulator of G protein signaling (RGS) domains accelerate the hydrolysis of GTP to inhibit G protein signaling. Some RGS domains are believed to be selective to specific Gα proteins although many are promiscuous towards several structurally-related related Gα proteins ^4–6^. The activities of many of these RGS domain proteins are not always known and many are regulated transcriptionally. Ultimately, ligands, receptors, and modulatory proteins control neuromodulator signaling and functions by the precisely controlled spatial and temporal presence at the site of the neurotransmission, leading to the defined control of behavior and physiological response to sensory stimulation.

Many optogenetic approaches modulate intracellular signaling, including GPCR and G-protein signaling. However, most of these approaches focused on strategies to activate cellular signaling pathways to initiate specific cellular events by the over-expression of exogenous proteins or engineered domains derived from endogenous proteins ^7–10^. The optogenetic inhibition of GPCR signaling is challenging to develop and inhibition requires the over-expression of proteins to target the endogenous enzymes or the expression of an exogenous protein that would either counteract or disrupt the endogenous signaling pathway ^11^. Other researchers attempted to inhibit G-protein signaling ^12, 13^, but the systems lacked rigorous testing of G-protein inhibition specificity and were only tested in cell culture. Disruptive approaches are essential to understand specific signaling pathways’ physiological consequences and timing in response to external stimuli.

In the current study, we use and modify the RGS2 domain to develop optogenetic approaches that selectively disrupt Gα_q/11_ by light-induced membrane recruitment. We name our system the **P**hoto-**i**nduced **m**odulation of **G**α protein – **I**nhibition of Gαq or PiGM-Iq. PiGM-Iq specifically inhibits Gα_q_ signaling. We test PiGM-Iq in *C. elegans*, *Drosophila*, zebrafish, and primary culture dorsal root ganglia neurons to alter mobility, learning and synaptic plasticity, and axon development, respectively. Additionally, we change the selectivity of PiGM by targeting Gα_i/o_ signaling using the RGS10 domain. Our newly created optogenetic tools are significant additions for studying cellular signaling in living organisms to understand complex biological functions and behavior. Future engineering will expand our method to selectively suppress other G protein signaling cascades to understand the higher-order processes of animals reacting to complex stimuli and environments.

## Results

### Membrane recruitment of minimal RGS2 domain to inhibit Gα_q_ signalling

To selectively inhibit Gα_q_ signalling associated with GPCR activation, our strategy is to locate the Gα_q_-suppressing RGS domain to its substrate optogenetically. Gα_q_ targeting the RGS domain would normally be found on the membrane where Gα_q_ is located or have such high affinity with GTP-bound Gα_q_ that it would be recruited to the membrane when Gα_q_ signalling is activated (Fig. 1A). Hence our strategy requires the identification of an RGS domain that fulfill the following criteria: 1) Can be mislocated to the cytosol; 2) does not have high affinity to GTP-bound Gα_q_ without optogenetic manipulation; 3) act selectively on Gα_q_ and not the structurally similar Gα_i/o_; and that 4) optogenetic recruitment of the RGS domain to the membrane is sufficient for activation. For the photodimerization system, we have chosen the CRY2PHR/CIBN photodimerizer pair because it recruits cytosolic fluorescent proteins to the membrane robustly with a high level of light sensitivity ^14^ (not shown).

**Figure 1.**
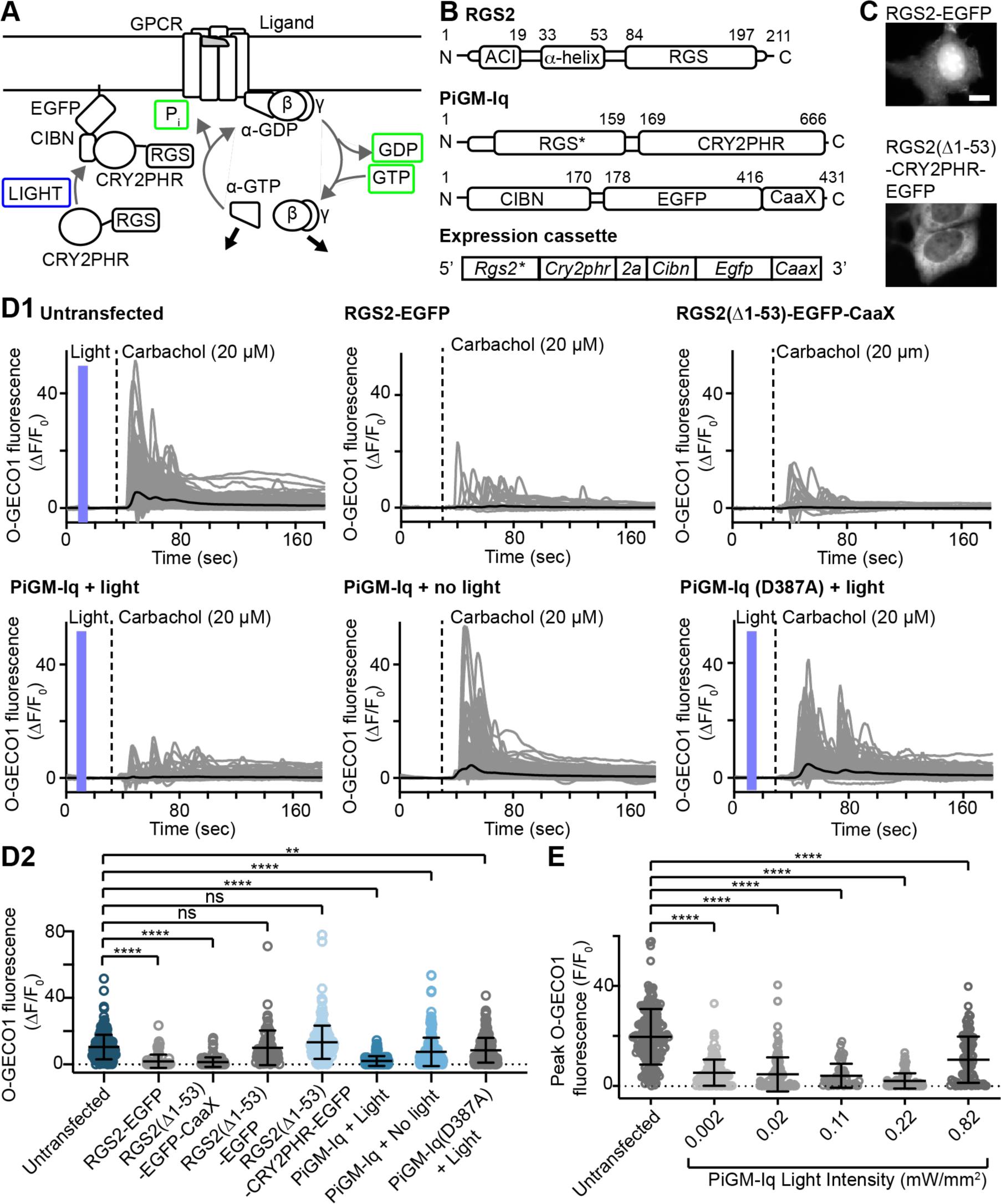
The design of the PiGM-Iq and basic characterization. (A) Proposed mechanism of PiGM-Iq in schematic form. Light-induced membrane recruitment of CRY2PHR and the associated RGS domain from RGS2 bring the RGS domain to the membrane substrate of GTP-bound Gα_q_ and catalyze the hydrolysis of GTP to GDP and the termination of Gα_q_ signaling. (B) The linear protein domain organization of wild-type RGS2 (top), the 2 components of PiGM-Iq (middle) and the recombinant DNA expression cassette (bottom) used in most of this study. Wild type RGS2 contains a N-terminal peptide that is reported to inhibit adenylate cyclase (denoted ACI – adenylate cyclase inhibitor), a transmembrane α-helix and a RGS domain at the C-terminal. In the PiGM-Iq design, The first 53 amino acids of RGS2 were removed and a N-terminus methionine was introduced to the remaining RGS domain was tethered to the photodimerization domain CRY2PHR. In some instances, mCherry was introduced at the C-terminus to visualize the localization of the protein. The second component of PiGM-Iq is the photodimerization domain CIBN anchored to the membrane via an EGFP with a palmitoylation CaaX sequence. For the expression of PiGM-Iq, the 2 components were co-expressed using a 2A bicistronic expression system. (C) Expression of C-terminal EGFP tagged RGS2 and RGS2 with truncated N-terminal tagged with CRY2PHR-EGFP in HEK293 cell. RGS2-EGFP is localized mostly in the nucleus whereas the RGS2 with N-terminal truncation and tethered to CRY2PHR-EGFP is in the cytosol. Scale bar is 10 µm. (D) Calcium elevation measured with O-GECO1 fluorescence in untransfected HEK293 cells and cells expressing different constructs in response to carbachol (20 µM) application. (D1) shows the time courses of calcium responses in individual cells (gray lines) and the mean responses (black lines), dotted black lines indicates the time of carbachol application and the blue boxes indicate time of blue light illumination. (D2) is the summary of the individual and mean peak amplitudes of the different conditions showed in (D1). (E) The effects of light intensity and PiGM-Iq at suppressing carbachol-induced calcium elevation with strongest effect seen at 0.22mW/mm^2^. For (D2) and (E), Kruskal-Wallis test and Dunn’s multiple comparison tests against untransfected sample were used and ** and **** indicate levels of significance of <0.005 and <0.0001 respectively. Graphs are shown as mean ± S.D.

To screen for the appropriate RGS domain, we applied the non-selective muscarinic receptor agonist carbachol to activate the endogenously expressed M3R receptor in HEK293 cells ^15^ and monitored the calcium elevation with the orange calcium indicator O-GECO1 ^16^. The calcium elevation induced by carbachol was associated with PLCß activation and attributed to intracellular calcium storage as expected (not shown).

We identified the RGS domain of RGS2 for our approach. The RGS domain of RGS2 has >8x selectivity to Gα_q_ over Gα_i_ ^3^. RGS2 contains an N-terminus that inhibits adenylate cyclase (amino acids 1-19) ^17–19^ and a membrane-targeted alpha helix (amino acids 33-53) ^20^. We eliminated the first 53 amino acids of RGS2 in our design to both remove the adenylate cyclase inhibiting peptide and membrane association region in our final design (Fig. 1B). Expression of the wild-type RGS2 with EGFP tag at the C-terminus (RGS2-EGFP, Fig. 1C) and a modified RGS domain from RGS2 artificially located to the membrane with CaaX motif (RGS2(Δ1-53)-EGFP-CaaX) strongly suppressed carbachol-induced calcium elevation (ΔF/F_0_ of 10.4 ± 7.4 in *n* = 255 untransfected cells vs. ΔF/F_0_ of 1.8 ± 4.0 in *n* = 102 RGS2-EGFP expressing cells and ΔF/F_0_ of 1.3 ± 2.9 in 140 RGS2(Δ1-53)-EGFP-CaaX-expressing cells), and that the expression of RGS2 without the first 53 amino acids fused to cytosolic EGFP failed to reduce calcium elevation (ΔF/F_0_ of 10.0 ± 10.4 in *n* = 80 cells), suggest the first 53 amino acids did not contribute to the ability of RGS domain from RGS2 to inhibit Gα_q_ function and that this segment is essential for its localization to the membrane to inhibit Gα_q_ (Fig. 1C & D). Interestingly, the wild-type RGS2 not only accumulates in the membrane but also strongly in the nucleus when expressed as an EGFP fusion (Fig. 1C). Optogenetic membrane recruitment of the truncated RGS domain from RGS2 with CRY2PHR/CIBN can completely abolish carbachol-associated calcium elevation in most expressing cells (ΔF/F_0_ of 2.0 ± 3.0, *n* = 243 cells) and the effect was reversed without light illumination (ΔF/F_0_ of 7.5 ± 8.6, *n* = 255 cells). The light-insensitive mutant CRY2PHR(D387A) lacked inhibition (ΔF/F_0_ of 8.5 ± 7.5, n = 255 cells). When we recruited the RGS domain to cytosolic-located CIBN, there was no significant change in calcium (ΔF/F_0_ of 13.1 ± 10.2, *n* = 217 cells)(Fig. 1D). However, it is worth noting there is a significant reduction of peak calcium responses without light and using the D387A mutant, suggesting there are some inhibitory effects associated with the background affinity of CRY2PHR to CIBN as previously reported ^21^ and/or the RGS2 domain still has some affinity to GTP-bound Gα_q_. Interestingly, only the C-terminal fusion but not the N-terminal fusion of CRY2PHR can achieve such strong and consistent inhibition of calcium increase (not shown). Another RGS domain from GRK2 is reported to have high selectivity over Gα_q_ ^22^, but the light-mediated recruitment of this RGS domain to the membrane did not suppress the carbachol-mediated calcium elevation as strongly as the RGS domain of RGS2. We also introduced two mutations (T137D, E141M) on the RGS domain of RGS2 based on the structure of GRK2 in an attempt to increase the selectivity of the RGS2 domain over Gα_i_ ^22, 23^ but this also failed to suppress the calcium increase as robustly as the minimal RGS2 in our original design (Suppl Fig. 1). Hence the truncated RGS domain of RGS2 remains the best protein domain for our approach. We named our approach ‘**P**hoto-**i**nduced **G**-protein **M**odulator – **I**nhibitor of Gα**_q_**‘ or PiGM-Iq.

### Characterisation of PiGM-Iq

Our modified RGS domain with CRY2PHR is located in the cytosol when fused with a C-terminal fluorescent protein, EGFP or mCherry (Fig. 1C) and is optogenetically recruited to the membrane repetitively and robustly (Suppl Fig. 2). The membrane recruitment of cytosolic mCherry achieved maximum efficiency with light intensity ∼0.22 mW/mm^2^, with higher light intensity achieving lower membrane recruitment. Using the calcium elevation associated with carbachol stimulation, the efficiency of PiGM-Iq also reached maximum efficiency ∼0.22 mW/mm^2^ with lower efficiency at higher light intensity (Fig. 1E). In terms of expression level, there is a weak correlation between the EGFP expression level in a 2A-based bicistronic expression cassette and the reduction of calcium elevation (ρ = −0.33), suggesting the effect does not require a high level of expression. There are also weak correlations in the absence of light with PiGM-Iq (ρ = −0.39) and with light-insensitive mutant D387A (ρ = −0.43) but not when the RGS domain of RGS2 is directly expressed as a CRY2PHR-EGFP cytosolic fusion protein, suggesting some background binding activity of CRY2PHR and CIBN contributes to a small degree of background RGS activity and the D387A mutant may still have some binding capacity to CIBN (Fig. 2A). As the selectivity of RGS2 towards Gα_q_ over Gα_i_ was reported as 8-fold previously ^3^, we were concerned that PiGM-Iq might act promiscuously to inhibit Gα_i_ and Gα_q_ signalling when overexpressed. To test the possible inhibitory effects of PiGM-Iq on Gα_i_, HEK293 cells were co-transfected with the GloSensor, a rat D2 dopamine receptor (D2R), and PiGM-Iq constructs. The GloSensor is a luciferase-based cAMP assay. When stimulated with isoprenaline, intracellular cAMP increases via activating the endogenous Gα_s_-coupled ß2 adrenergic receptor expressed in HEK293 cells.

**Figure 2.**
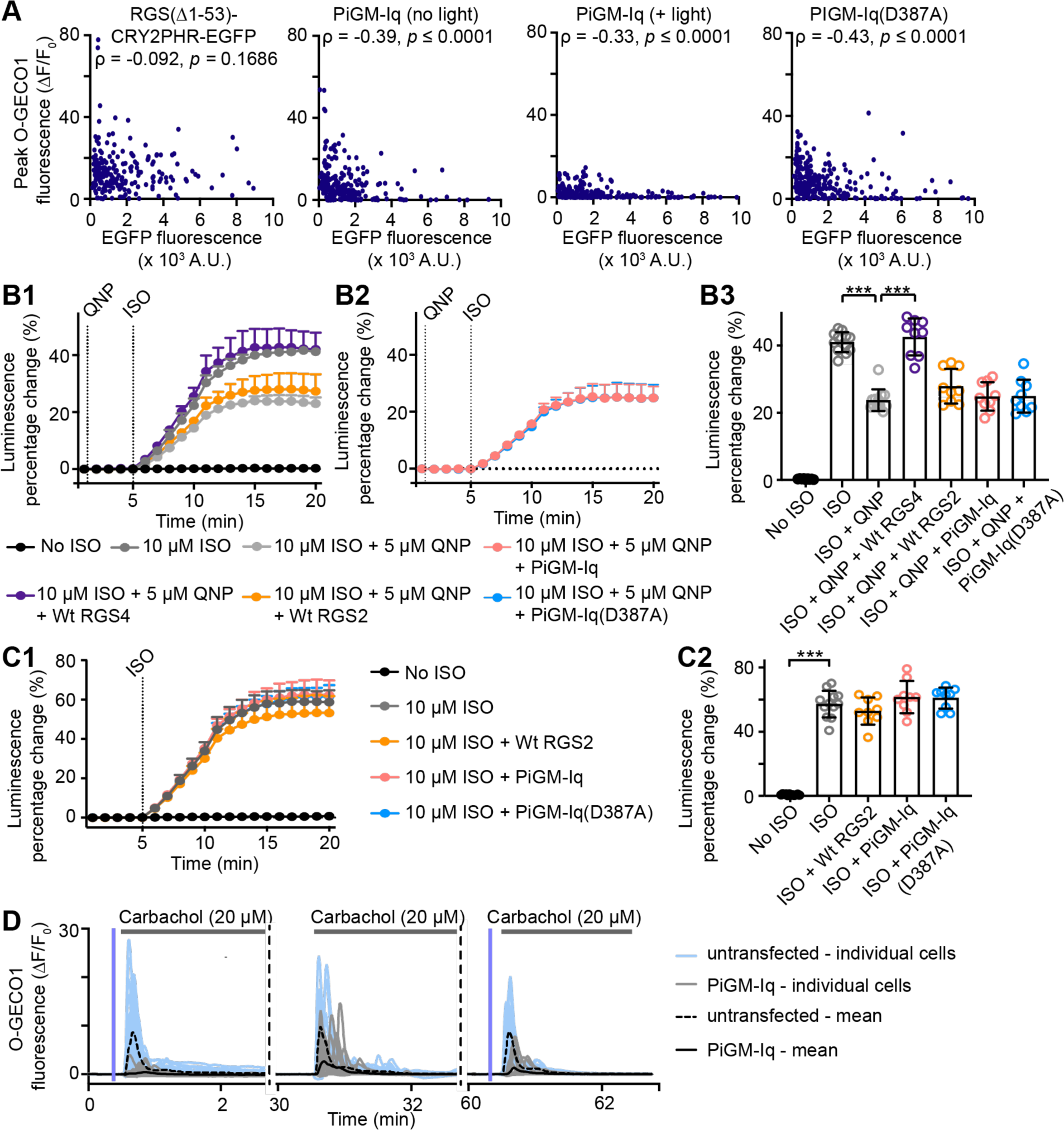
The characterization of the PiGM-Iq properties. (A) The correlation of carbachol-induced O-GECO1 fluorescence changes (ΔF/F_0_) and EGFP fluorescence in cells expressing RGS(Δ1-53) fused to CRY2PHR-EGFP (far left), PiGM-Iq bicistronic expression cassette with no light illumination (middle left), PiGM-Iq expression cassette with light illumination (middle right) and PiGM-Iq with D387A mutation in CRY2PHR (far right). With the exception of RGS(Δ1-53)-CRY2PHR-EGFP, significant but weak negative correlations were detected between calcium elevation and EGFP expression with Spearman correlation analysis. (B1) The luminescence folds change associated with isoprenaline (ISO)-induced cAMP increase in HEK293 cells exogenously expressing GloSensor luciferase and D2R dopamine receptor in different conditions. Isoprenaline (ISO) elevates cAMP and luciferase activity via ß2-adrenergic receptor activation and Quinpirole suppressed the cAMP elevation via D2R activation of Gα_i_. Wild-type RGS4 was able to disrupt the effects QNP whereas wild-type RGS2, PiGM-Iq and PiGM-Iq(D387A) did not disrupt the effects of QNP. Light illumination was applied to all samples. (B2) The summary of the final peak luminescence folds changes. *** indicates adjusted P-value of 0.0005 compared to ISO + QNP sample. (C1) The luminescence folds change associated with isoprenaline (ISO)-induced cAMP increase in HEK293 cells with light stimulation. Expression of wild-type RGS2, PiGM-Iq and PiGM-Iq(D387A). *** indicates adjusted P-value of 0.0003 compared to ISO sample. Kruskal-Wallis test followed by Dunn’s multiple comparisons tests were used to detect the difference between the indicated sample and ISO + QNP sample in (B) and the ISO sample in (C). (D) The individual and mean calcium-elevation in HEK293 cells measured with O-GECO1 after light-induced PiGM-Iq activation. After 30 min washout of carbachol and recovery of the initial inhibition, carbachol application was able to evoke calcium elevation in PiGM-Iq-expressing cells when light was omitted. After another 30 min, light illumination of PiGM-Iq cells was able to reduce carbachol induced calcium elevation. Light illumination was indicated by vertical blue bars prior to carbachol application. Graphs are shown as mean ± S.D.

Co-application of isoprenaline and D2R agonist quinpirole to activate Gα_i_ signaling leads to the suppression of cAMP production (23.76 ± 3.24%, *n* = 12 cells vs. 41.02 ± 2.97%, *n* = 12 cells for isoprenaline only; Fig. 2B). Light activation of PiGM-Iq did not inhibit the inhibitory effects of quinpirole on isoprenaline induced cAMP increase (24.88 ± 4.22, *n* = 9 cells; Fig. 2B). Expression of wild-type RGS2 (27.91± 5.14%, *n* = 9 cells) and PIGM-Iq(D387A) with light illumination (24.95 ± 4.92%, *n* = 9 cells) also have no change on the quinpirole-D2R effects but the expression of wild-type RGS4 (42.58 ± 4.50%, *n* = 9 cells) inhibited the suppression effect of quinpirole. We also did not observe any inhibitory effects of PiGM-Iq on ß2 adrenergic receptor/Gα_s_ induced elevation of intracellular cAMP (61.56 ± 15.05%, *n* = 9 cells vs. 57.23 ± 8.29%, *n* = 12 cells with isoprenaline only; Fig. 2C). These results suggest minimal cross-reactivity of PiGM-Iq with either Gα_s_ or Gα_i_ and verify selective inhibition of Gα_q_.

To test whether the PiGM-Iq effect is reversible, untransfected and PiGM-Iq-expressing cells were challenged with carbachol at three time points with 30-minute intervals, and the light was only applied during the first and third carbachol application. Most PiGM-Iq-expressing cells responded to carbachol as expected with calcium elevation during the second carbachol application without light stimulation (ΔF/ F_0_ = 5.98 ± 3.98, *n* = 29 cells). A small subset of cells failed to respond to carbachol during the second ‘light-less’ stimulation (defined as ΔF/ F_0_ < 1.0, *n* = 3 out of 29 total cells) (Fig. 2D). In PiGM-Iq expressing cells, light stimulation strongly suppressed the first carbachol challenge (ΔF/ F_0_ = 0.88 ± 1.08 with 20/29 cells with ΔF/ F_0_ < 1.0). The third carbachol stimulation is slightly less effective, but light stimulation suppressed the response of PiGM-Iq expressing cells (ΔF/ F_0_ = 1.97 ± 2.49 with 16 out of 29 total cells with ΔF/ F_0_ < 1.0). In non-expressing control cells, the 3 carbachol challenges with the same light stimulation pattern resulted in ΔF/ F_0_ of 12.54 ± 3.68, 13.60 ± 4.47, and 11.09 ± 4.2 (*n* = 29 cells) with 1/29, 0/29 and 3/29 cells failed to respond to carbachol stimulation respectively. Just to note that the calcium response of PiGM-Iq-expressing cells in light-less carbachol challenge is significantly smaller than non-expressing cells (P < 0.0001, unpaired t-test). Our experiments suggest the PiGM-Iq system is reversible, but there may be some residual inhibition that is enhanced with high expression of proteins (unquantified observation).

### Activation of PiGM-Iq reduces *C. elegans* mobility

The *C. elegans* Gα_q_ homologue EGL-30 shares 82.8% sequence homology with the mammalian version of the protein (Suppl material 2). Hence we predicted that PiGM-Iq could efficiently suppress EGL-30 activation. EGL-30 complete loss-of-function mutants are not viable due to developmental problems, but EGL-30 mutant lines with reduced EGL-30 activities have uncoordinated, slow movement and reduced pharyngeal pumping ^24^. The reduced mobility is associated with the decreased acetylcholine release at the neuromuscular junction with reduced Egl-30 activity ^25^(Fig. 3A). We were able to generate viable PiGM-Iq and PiGM-Iq(D387A) dark mutant expressing worms using a pan-neuronal *snb-1* promoter. The EGFP fluorescence associated with the membrane-tethered CIBN is observable in *C. elegans* neurons (Fig. 3B). We could not generate animals expressing wild-type mammalian RGS2 as these animals were not viable. Although not quantified, we observed that worms expressing a high level of PiGM-Iq and PiGM-Iq(D387A) grew slower or sometimes reached developmental arrest. Slowed development suggests possible background activities with modified RGS2 domain when expressed at high intracellular concentration and possible background binding of CRY2PHR and CIBN. The worms with a lower level of PiGM-Iq expression, as visualised with EGFP fluorescence, do not have obvious developmental defects or reduced viability.

**Figure 3.**
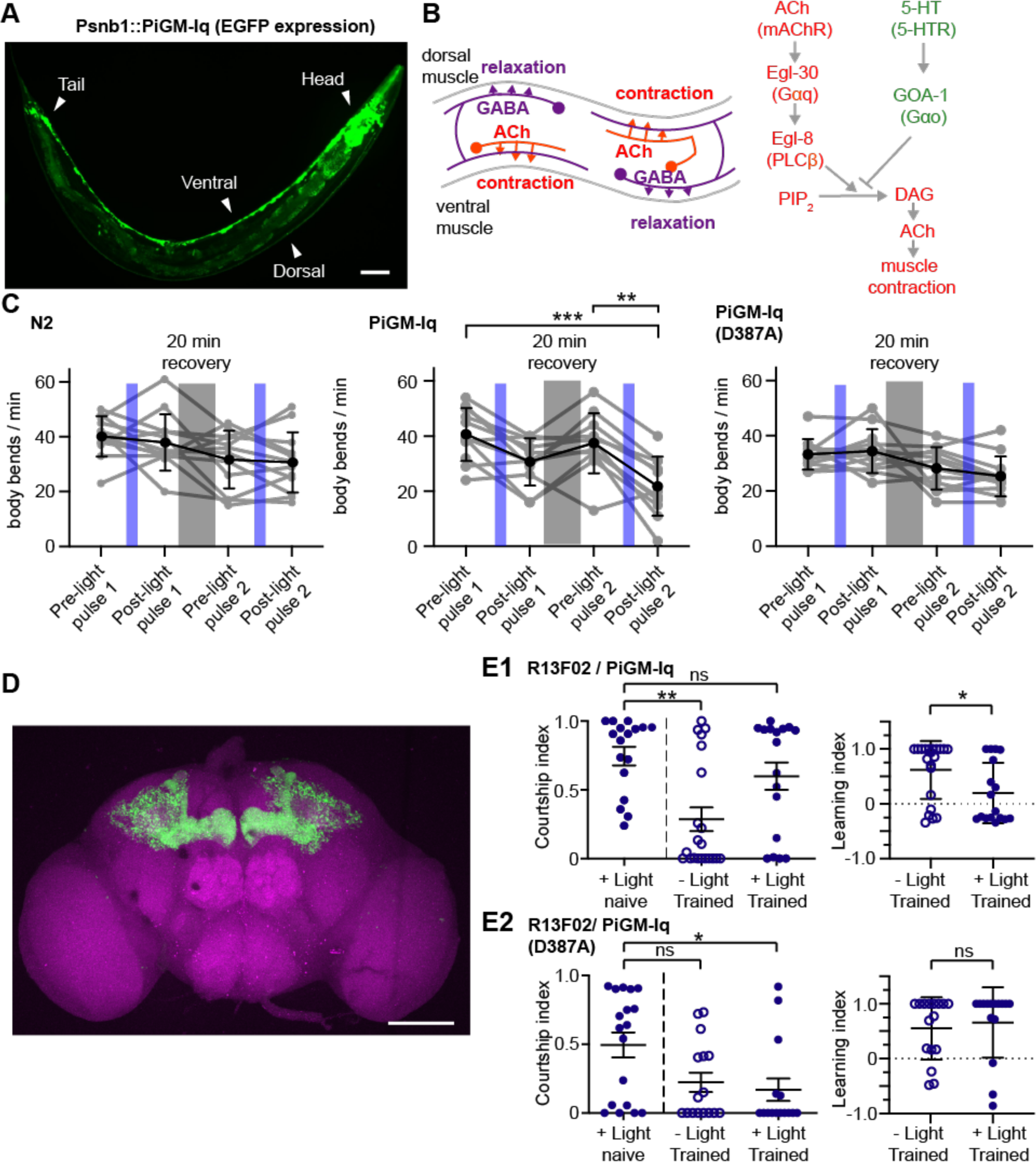
PiGM-Iq activation leads to reduced mobility in *C. elegans* and disrupts courtship learning in *Drosophila*. (A) Schematic of the neurocircuitry coordinates movements of dorsal and ventral body walls. Cholinergic innervation via the activation of Egl-30/Gαq-linked muscarinic receptor leads to muscle contraction. Disruption of Egl-30 leads to the disruption of movement coordination and reduced mobility (modified from ^25, 48^). (B) Expression of EGFP along the ventral cord of *C. elegans* is consistent with pan-neuronal expression of PiGM-Iq cassette. (C) Light illumination in PiGM-Iq-expressing worms lead to reduced mobility not seen in wild-type/N2 and PiGM-Iq(D387A) expressing worms. (D) The expression pattern of EGFP in the brain of R57C10/PiGM-Iq flies showing mushroom body targeted expression. The synapses in the brain are labelled with anti-Bruchpilot (Brp) antibodies and additionally stained with secondary antibody with Alexa-594. (E) The courtship and learning indices of R57C10/PiGM-Iq (E1) and R57C10/PiGM-Iq(D387A) (E2) for courtship learning. In naïve male flies expressing PiGM-Iq or PiGM-Iq(D387A), light illumination does not alter the attempted courtship behavior (left). In PiGM-Iq-expressing flies without light illumination and D387A flies with and without light illumination, the male flies exhibited reduced courtship behavior after conditioning with mated female, and this were also reflected as increased learning indices. In PiGM-Iq-expressing male flies, light illumination prior to conditioning with mated female disrupted learning of the conditioning and the male flies still exhibited significant courtship towards mated female. For (C) ** and *** indicate significance of P = 0.0038 and P = 0.0004, respectively. The significance in (C) calculated with ordinary one-way ANOVA followed by Tukey’s multiple comparison tests between all treatment groups. Courtship Indices in (E) were tested with Kruskal-Wallis test followed by Dunn’s test against naïve flies. Learning indices were tested using Mann-Whitney test (E). * and ** indicate P < 0.05 and P < 0.01, respectively. Graphs are shown as mean ± S.D. Scale bars in (D): 100 µm.

We measure the movements of wild-type, PiGM-Iq, and PiGM-Iq(D387A) worms in response to 2 pulses of blue light illumination. There was no significant reduction of body bends per minute in wild-type or PiGM-Iq(D387A) worms in response to the first or second pulses of blue light (For values, see Suppl table 1; Fig. 3C). However, the PiGM-Iq worms showed a statistically insignificant reduction in movement after the first blue light pulse and showed a statistically significant decrease after the second pulse (For values see Suppl table 1). The background locomotion of the 3 lines was not significantly different before light illumination. Our results are consistent with the expected reduced mobility seen with the EGL-30 mutants ^25^.

### LexA-inducible PiGM-Iq activation suppresses *Drosophila* courtship learning

In *Drosophila melanogaster*, Gα_q_ homologue shares 84.7% homology to the mammalian Gα_q_ (suppl material 3). Therefore, we predicted PiGM-Iq based on mammalian RGS2 would work in flies. We generated PiGM-Iq lines inducible through the LexA/LexAop bipartite expression system ^26, 27^ allowing these tools to be expressed cell-type specifically using the vast array of LexA driver lines.

In insects, octopamine is a neurotransmitter and neuromodulator that shares a similar structure to noradrenaline, a mammalian ligand. In the mushroom body, the insect center of olfactory learning, octopamine specifically acts through the Gα_q_ coupled OAMB-AS receptor ^28^, making it distinct from other octopamine and dopamine receptors in the fly that works through Gα_s_. Significantly, the OAMB receptor in the mushroom body was previously shown to be necessary for courtship conditioning ^29^, a process through which male flies are less likely to court a female fly after a previous courtship rejection. We thus hypothesized that the inactivation of Gα_q_ signalling via PiGM-Iq should suppress this behaviour. In previous optogenetic work with channelrhodopsins in *Drosophila*, blue light illumination has been avoided owing to its poor penetration of the melanised adult cuticle ^30^. However, the ventral portion of the fly head capsule is translucent and lacks pigment, and we overcame this problem by illuminating flies from the ventral side (Suppl Fig. 3A).

In wild-type control flies, the male courtship response was suppressed as expected after pre-conditioning with non-receptive (mated) female flies (Suppl Fig. 3, Suppl tables 2 & 3), and illumination with the blue light period during the pre-conditioning period did not alter the courtship conditioning response (Suppl Fig. 3B). Flies expressing PiGM-Iq either solely in the mushroom body (driven by the MB-specific R13F02-LexA driver, Fig. 3D) or pan-neuronally (via the pan-neuronal R57C10-LexA driver) similarly exhibited courtship conditioning without exposure to blue light (Fig. 3E, Suppl Fig. 3D & Suppl table 2). However, the courtship conditioning response was suppressed upon blue light exposure, and flies continued to court females even after previous rejections (Fig. 3E1, Suppl Fig 3D, Suppl table 2). Importantly, a disruption of learning was not observed in flies expressing the light-insensitive mutant version of PiGM-Iq (CRY2PHR with D487A mutation; Fig. 3E2, Suppl Fig. 3E; Suppl table 2 & 3). Our data show that the PiGM system is light-specific when induced with the GAL4 system and can be used to disrupt specific signalling pathways in neuronal subtypes.

### Light activation of PiGM-Iq alters serotonin-evoked axon growth cone turning in cultured DRG neurons

In our previous study, we showed the activation of metabotropic serotonin receptors in growth cones of cultured DRG neurons directed axon turning depending on the activated receptor ^31^ (Fig. 4A). A 100 μM serotonin gradient resulted in repulsion of growth cones mediated by the Gα_i_-coupled 5-HT1b receptor, whereas 50 μM of serotonin gradient resulted in attractive turning mediated by the Gα_q_-coupled 5-HT2a receptor. Attractive turning was observed when untransfected growth cones were exposed to blue light and 50 μM of serotonin (11.28 ± 4.34°, *n* = 12 cells; Fig. 5B). Asymmetric PiGM-Iq activation (6x 2 sec pulses, 5 min interval; Fig. 4A) of growth cones in the direction of a 50 μM serotonin gradient resulted in repulsive turning (−7.12 ± 4.51°, *n* = 13 cells), compared to PiGM-Iq-expressing growth cones exposed to 50 μM of serotonin without light illumination that maintained attractive turning (11.03 ± 3.24°, *n* = 11 cells; Fig. 4B).

**Figure 4.**
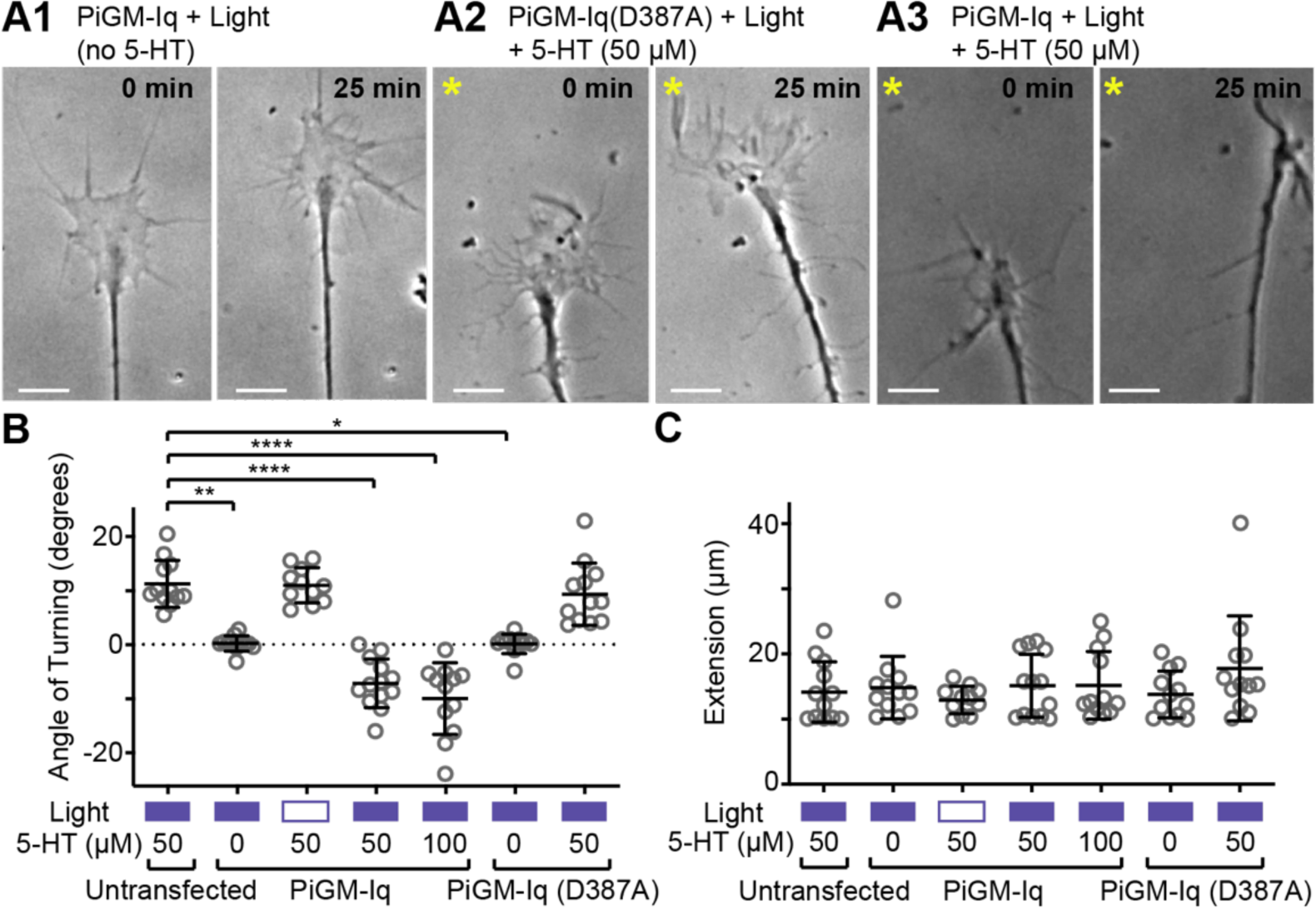
PiGM-Iq activation alters the effects of serotonin on axon growth cone turning. (A) Examples of growth cones of PiGM-Iq-expressing DRG primary neurons in response to light and no serotonin (5-HT) application (A1) and 50 µM 5-HT from the top left of the growth cone (indicated by the asterisk)(A2 and A3). In (A2), the DRG neuron expressed the light insensitive D387A mutant of PiGM-Iq and 5-HT application leads to attractive turning. In (A3), the light activation of the non-mutant version of PiGM-Iq leads to repulsive turning to 5-HT. Blue light illumination is restricted to the side of serotonin application using a DMD device coupled to the microscope. (B) Summary graphs of number of growth cone in response to light (indicated by blue solid box), no light (indicated by empty box) and the concentration of 5-HT as indicated. In the absence of 5-HT, PiGM-Iq and PiGM-Iq(D387A)-expressing growth cones exhibit no orientation preference and straight projection in the presence of light. In the presence of 50 µM 5-HT, untransfected growth cones with light stimulation, PiGM-Iq-expressing growth cones with no light stimulation and PiGM-Iq(D387A)-expressing growth cone with light stimulation turns towards the 5-HT source. In PiGM-Iq-expressing growth cones with light stimulation and 50 and 100 µM 5-HT, the growth cones turn away from the 5-HT source. (C) Light stimulation and PiGM-Iq activation have no significant effects on the length of axon extension. Statistical significances were tested with Kruskal-Wallis test followed by Dunn’s test against untransfected subject to 50 µM serotonin treatment if significance was detected. *, **, **** indicate significance of P < 0.05, 0.01 and 0.0001, respectively. Scale bars in (A): 5 µm. Graphs are shown as mean ± S.D.

**Figure 5.**
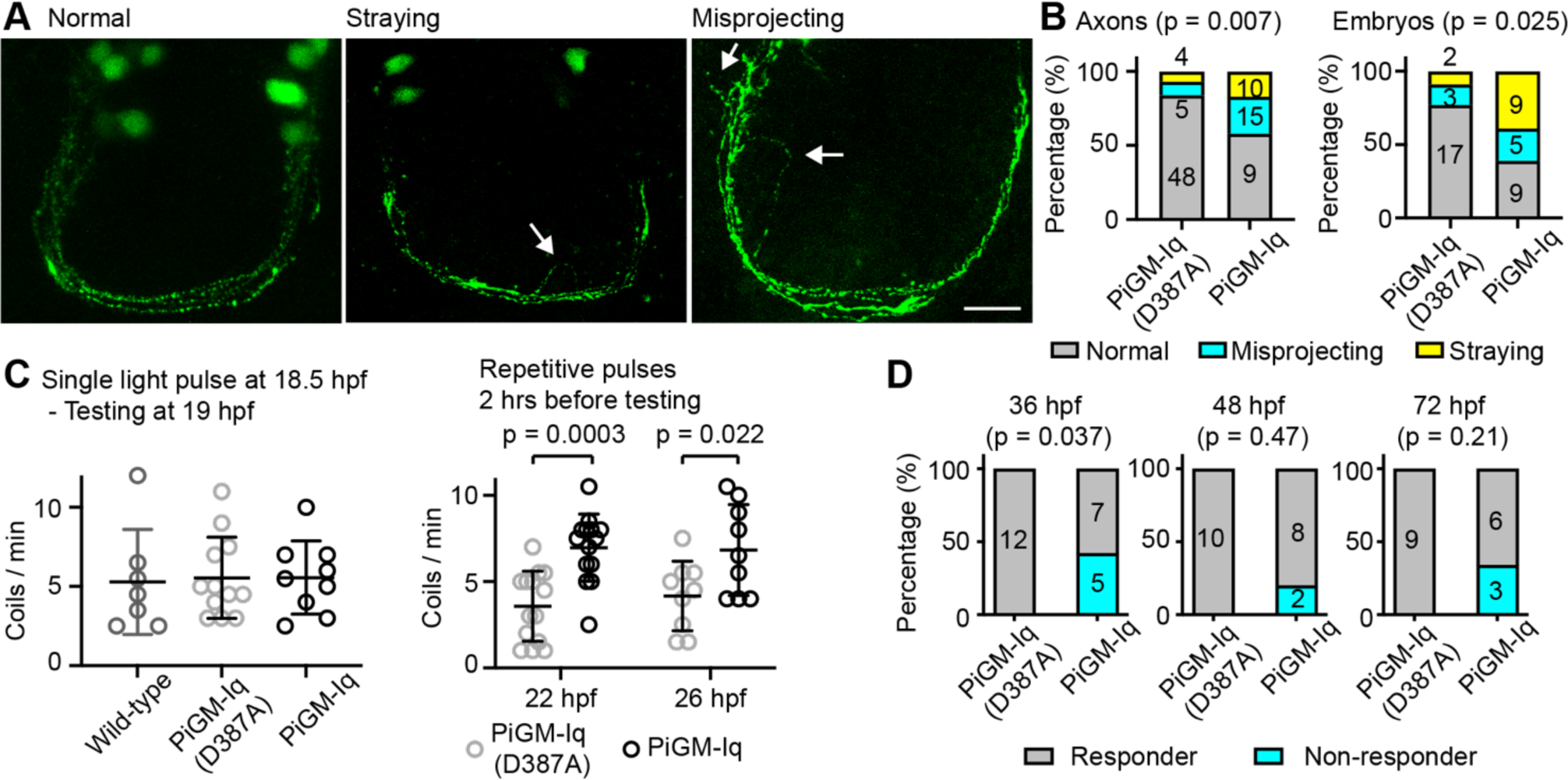
Disruption of zebrafish axon development and the functional effects of PiGM-Iq. (A) Example images of axons in zebrafish embryos showing what were classified as ‘straying’ (middle) and misprojecting (right) compared to normal (left). The regions of abnormalities were indicated by the arrows. Summary graph and significance of the number of axons (left) and embryos (right) at 26 hpf with described abnormalities in PiGM-Iq(D387A) and PiGM-Iq-expressing zebrafish embryos in response to repetitive light treatment. (C) Graphs summarizing the number of spontaneous coiling measured in the different strains in the described treatments in wild-type, PiGM-Iq and PiGM-Iq(D387A) animals. (D) Graphs summarizing the number of PiGM-Iq(D387A) and PiGM-Iq animals that respond to touch at the different time points. There is a greater proportion of animals no responding to touch in the PiGM-Iq-expressing animals at all 3 time points tested although only the 36hpf stage is statically significant. Significances in (B) were tested with Chi-square using the absolute numbers, significance in (C1) were tested with Kruskal-Wallis test, significance in (C2) were tested with Sidak’s multiple comparison test and significance in (D) were tested with Fisher’s exact test using the exact numbers. * and *** indicate indicate P = 0.0222 and P = 0.0003, respectively in (C2). Graphs in (C) are shown as mean ± S.D.

This change of attractive turning to repulsive turning was not observed when the light-insensitive D387A mutant version of PiGM-Iq (Fig. 4A2) was used in the presence of serotonin and light stimulation (9.39 ± 5.76 °, *n* = 12 cells; Fig. 4B). Repulsive turning has been observed in response to 100 μM serotonin exposure in untransfected growth cones ^31^, and was also observed with PiGM-Iq-expressing growth cones in response to light stimulation and 100 μM serotonin exposure (−9.94 ± 6.63°, *n* = 12 cells). In the absence of serotonin, PiGM-Iq-expressing growth cones maintained straight growth trajectories with asymmetric light stimulation (0.26 ± 1.42°, *n* = 12; Fig. 4A & B), similar to that observed with growth cones expressing the light-insensitive PiGM-Iq(D387A) stimulated with blue light and in the absence of serotonin (0.19 ± 1.81°, *n* = 12; Fig. 5A & B). Light illumination had no effects on the growth cone’s extension rate (Fig. 4C). Overall, these results suggest PiGM-Iq activation is sufficient to suppress Gα_q_-signaling associated with 5-HT2a receptor activation leading to disrupted attractive turning but does not disrupt Gα_i_-signaling associated with 5-HT1b receptor.

### Light activation of PiGM-Iq disrupts axon development

In many species, serotoninergic neurons are among the earliest to emerge in the developing brain. Ventral diencephalic serotonergic neurons are the first to emerge in the developing brain of zebrafish embryos and send short-range projections contralaterally ^32^. Given the early development of ventral diencephalic neurons and the sequence homology between *Danio rerio* and mammalian Gα_q_ proteins (Suppl material 4), we choose the circuit to investigate the effect of PIGM-Iq-induced Gα_q_ inhibition on axon guidance.

PiGM-Iq or the light insensitive PiGM-Iq(D387A) was introduced into zebrafish using a Tol2-transposase system to limit expression to *Tph2* serotonergic neurons using Gal4-UAS line ^33^. The membrane-bound EGFP of the PiGM-Iq expression cassette is observed in the ventral diencephalic (Fig. 5A), the hindbrain, the dorsal raphe serotonergic neurons and the spinal cord ^33^. Blue light stimulation (24 x 10 s pulse at 5 min interval, applied over 2 hours) of PiGM-Iq expressing zebrafish embryos at 24-26 hours post fertilisation (hpf) leads to increased numbers of PiGM-Iq-expressing axons ‘straying’ and ‘misprojecting’ (Fig. 5A & B) compared to PiGM-Iq(D387A)-expressing axons (p = 0.007, Chi-square test) and increased fraction of PiGM-Iq-expressing embryos with ‘straying’ or ‘misprojecting’ axons (p = 0.025, Chi-square test) when imaged at 26 hpf (Fig. 5A & B). Interestingly, a single pulse of light stimulation did not alter axon development (not shown).

We applied light stimulation during development to test whether PiGM-Iq changed the development of spontaneous coiling and startle response associated with disruption of axon guidance in the zebrafish. The spontaneous coiling (Fig. 5C) and touch-startle response (Fig. 5D) were measured in light-treated embryos. With spontaneous coiling, a single 60 s light pulse at 18.5 hpf did not alter the spontaneous coiling rate at 19hpf in PiGM-Iq expressing embryos compared to wild-type and the light insensitive PiGM-Iq(D387A) (Fig. 5C). Repetitive blue light stimulation (24 x 10 s pulses with 5 min interval) at 20 and 24 hpf resulted in an increase in spontaneous coiling at 22 and 26 hpf, respectively (Fig. 5C). With touch-startle-response, light stimulation (24 x 10 s pulses with 5 min interval) at 34, 46 and 70 hpf result in 5 out of 12, 2 out of 10 and 3 out of 9 embryos not responding to touch at 36, 48 and 72 hpf respectively (Fig. 6D). In light insensitive PiGM-Iq(D387A) embryos, similar light stimulation did not result in embryos not responding to touch at the same corresponding development stages (p = 0.0373 for 36 hfp, non-significant for 48 and 72 hpf, Fisher’s exact test; Fig. 5D).

**Figure 6.**
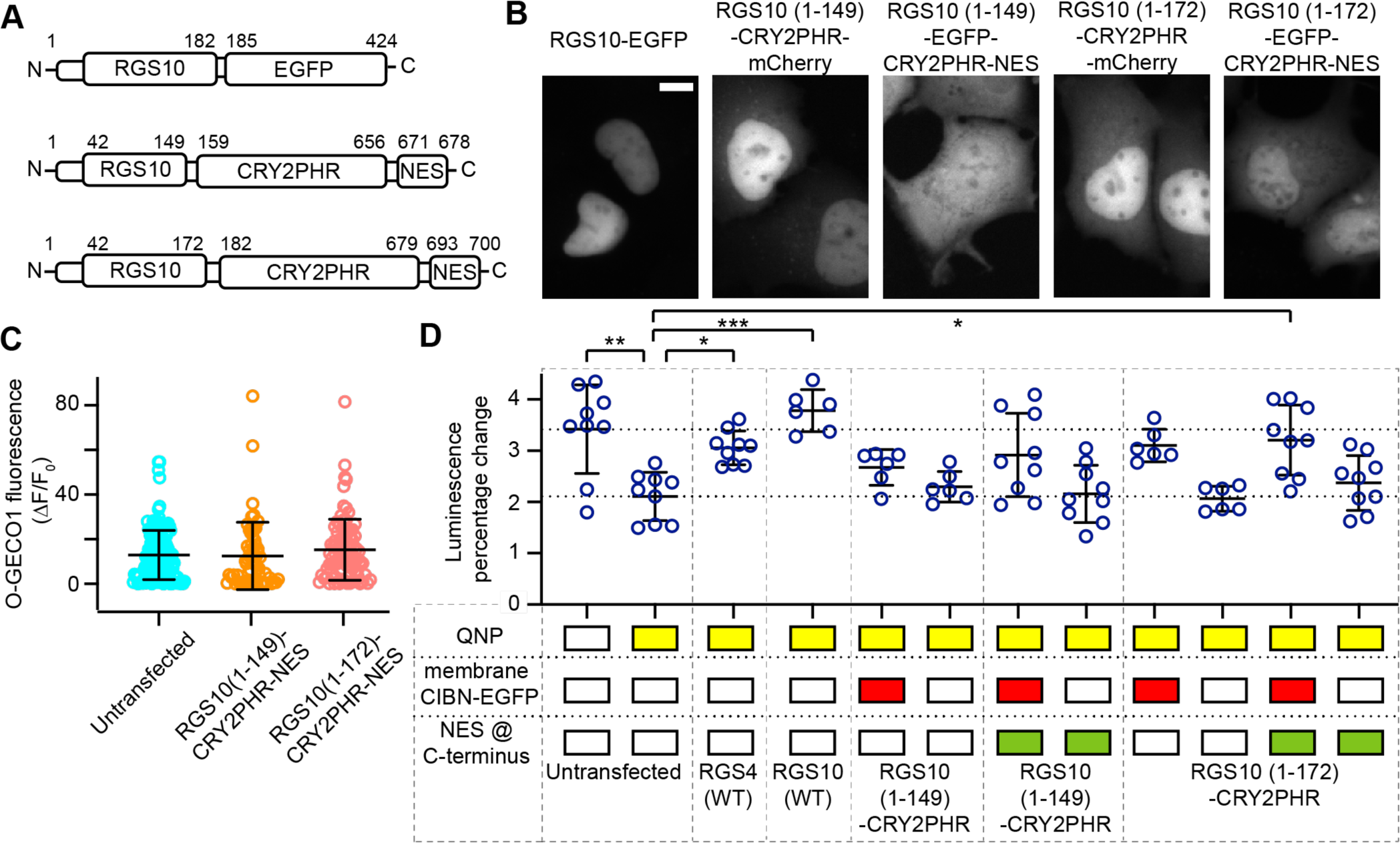
Alteration of the approach for optogenetic inhibition of Gα_i_ signaling. (A) Schematic representation of the RGS10 structure and the modification for the PiGM-Ii design. (B) representative EGFP images of wild-type RGS10 directly tethered to EGFP, truncated RGS10 (1-149 and 1-172) to CRY2PHR-mCherry and EGFP-CRY2PHR-NES. RGS10 has a dominant nuclear localization and the truncation and introduction of NES reduced the nuclear localisation of the RGS domain although in the cases of RGS10 (1-172) the expression is still predominantly nuclear with some cytosolic expression. (C) The responses to carbachol-induced calcium increase in HEK293 cells when expressing PiGM-Ii with truncated RGS10 with NES localization sequence. (D) The GloSensor luciferase assay result summary of cells expressing different constructs to quinpirole-D2 dopamine receptor inhibition of ß2-adrenergic receptor-cAMP increase. Expression of wild-type RGS4 and RGS10 can both suppress quinpirole-induced suppression of cAMP-Glosensor signal. Both truncated RGS10 (1-149) and RGS10 (1-172) with NES can also achieve the suppression of quinpirole response. HEK293 cells in all conditions were exposed to light stimulation and isoprenaline stimulation. Results in (C) were not significant and *, **, *** in (D) indicated significance levels of < 0.05, < 0.01 and < 0.001 respectively with Kuskal-Wallis test followed by Dunn’s multiple comparison tests against untransfected cells with quinpirole stimulation.

### Modified RGS10 domain for the selective disruption of Gα_i_ signaling

Based on the success of PiGM-Iq, we believe the same strategy will allow selective inhibition of other Gα proteins and we tested disruption of Gαi signaling using a Gα_i_ selective RGS domain. A previous study used the RGS domain of RGS4 to disrupt Gα_i_ signaling ^13^. However, RGS4 is known to interact with Gα_q_ and Gα_i_. When the RGS domain of rat RGS4 was used in the PiGM-I system, we observed a partial inhibition of muscarinic receptor-mediated calcium increase in our initial screen, consistent with the observation that the RGS domain of RGS4 can interact with Gα_q_ as well as Gα_i_. We selected the RGS domain from RGS10, which was reported to have a higher selectivity to Gα_i_ ^34^. Wild-type RGS10 tagged with EGFP showed exclusive nucleus localization. When the N-terminal segments of RGS10 (amino acids 1-149 or 1-172) were expressed as CRY2PHR-mCherry fusion protein, there was an accumulation of the fusion protein in the nucleus as well. Incorporating a nuclear export sequence (NES) could re-direct the RGS(1-149) domain to the cytosol so that cytosolic and nuclear fluorescence was observed. The NES reduced but did not eliminate the RGS10(1-172) accumulation in the nucleus (Fig. 6). Further truncation of the RGS10 domain to the N-terminal 29 amino acids excluded the CRY2PHR-mCherry-NES from the nucleus completely (not shown).

The C-terminus-truncated RGS domain of RGS10 did not significantly suppress carbachol-induced intracellular calcium increase when recruited to the membrane with light (Fig. 6C). Using the GloSensor cAMP-dependent luminescence assay, light-induced membrane recruitment of both C-terminus truncated RGS domain from RGS10 (1-145 and 1-172) disrupts the dopamine D2 mediated suppression of isoprenaline evoked cAMP increase, suggesting that Gα_i_ signaling can be disrupted by the RGS domain from RGS10 using the similar strategy as PiGM-Iq (Fig. 6D). We named this approach Photo-induced G protein Modulation – Inhibition Gα_i_ or PiGM-Ii.

## Discussion

With advances in optogenetic and chemogenetic tools, there is an abundance of activation tools associated with G-protein signaling pathways with unique and distinct properties. Inhibitory tools to selectively disrupt G-protein signaling are challenging to develop and rarely selective for a single Gα protein. Our current study described and characterized two optogenetic approaches, PiGM-Iq and PiGM-Ii, that inhibit Gα protein signaling using light-mediated membrane recruitment of the modified RGS domain. While the designs were identical to that previously published ^12, 13^, we have extensively characterized the approaches and focused on limiting the cross-reactivity of the approaches by modifying the RGS domain used in the study. We have also demonstrated the utilities of the approaches in behaving animal models of *C. elegans*, *Drosophila*, and zebrafish in addition to primary rat neuronal culture. In these models, PiGM-Iq was highly effective and efficient at disrupting the cellular and organismic responses associated with Gα_q_ signalling. Although not shown in the current study, we can adapt our PiGM-I approaches with chemical dimerization systems ^35, 36^ to develop the corresponding chemogenetic versions of the tools. Our observations indicated there are some background activities associated with PiGM-Iq when expressed at a high level, which can be the result of background CRYPHR/CIBN in the dark and/or the affinity of the RGS domain towards activated GTP-bound Gα. It is possible to reduce the background activity and effect duration using newer versions of CRY2/CIB1 mutants ^21^ or utilize different photodimerization systems ^37, 38^. However, at its current state, a lower expression level appears to reduce the background activity to a negligible amount.

In our study, we did not observe significant undesired cross-reactivity of RGS domains towards Gα_q_ and Gα_i_ based on our GloSensor, calcium imaging, and axon turning experiments. However, such non-selective activity can’t be ruled out with the sensitivity and consistency of these assays. Again, reducing the expression level may be the simplest solution to limit the effects of Gα cross-reactivity. Interestingly, we also tested the RGS domain from SNX13 that was previously reported to be Gα_s_-selective ^39^ but we did not observe any effects on cAMP generation upon stimulation (not shown).

Compared to channelrhodopsins, the light sensitivity of CRY2/CIB1 photodimerization is much higher ^40^, and high levels of expression level and increased light intensity do not correlate to better performance of PiGM-I. On the contrary, high levels of expression often increase the basal activities of the tools, and the membrane recruitment becomes less efficient when light intensity exceeds 0.22 mW/mm^2^. This high level of light sensitivity translates to easier instrumentation for *Drosophila* experiments where blue light achieves stimulation through intact cuticles.

One of the main drawbacks of many optogenetic and chemogenetic tools to activate and inhibit cellular signaling is that they lead to a non-discriminative activation or inhibition of G-protein signaling of the whole cell unless the activation is optically controlled to a specific cellular location, and this may not be possible freely moving animals *in vivo*. The unmodified form of our PiGM-I would also simultaneously disrupt all the receptors activating the same Gα_q_ or Gα_i_ in the cell without discriminating different receptors. At its current version, PiGM-Iq should be used as a complementary tool to siRNA or CRISPR-mediated knockdown if investigating the roles of specific receptors. It is theoretically possible to achieve receptor-specific optogenetic disruption using the active receptor-specific nanobodies ^41, 42^ to restrict the PiGM-I effects to the proximity of the activated receptor. Proximity achieves a more specific disruption of G-protein signaling to the receptor of interest. Although theoretically possible, this may still have limitations in the temporal precision as binding of nanobodies to activated GPCR tends to have a slower time scale, and heterodimeric GPCR or GPCR co-localization has been reported. Nevertheless, this approach could warrant future efforts in the development of the PiGM-I approach.

## Methods

### Molecular cloning and plasmid generation

The plasmids and constructs were generated using standard molecular cloning techniques with appropriate digestion enzymes. Overlap extension polymerase chain reaction was used to generate some fusion proteins and introduce mutation in the recombinant DNA. For experiments in HEK293 cells, the expression cassettes were inserted in the pcDNA3 or pcDNA3.1+ (hygromycin) expression vectors (Thermo Fisher Scientific, MA, USA). For generation of stable cell lines, a second-generation lentivirus transfer vector with a CMV promoter was used. The coding sequences of RGS domains were extracted rat brain cDNA using PCR and primers designed based on sequences in NCBI database (RGS2 NCBI reference: NM_053453.2, RGS10 NCBI reference: NM_019337.1). D2R was also extracted from rat brain cDNA library (NM_001409379.1). O-GECO1-expression vector (#46025) was acquired from Addgene and subcloned into the pLenti-CMV transfer vector. GloSensor plasmid was acquired from Promega (WI, USA). Mammalian codon-optimized CRY2PHR and CIBN sequences were synthesized by Genscript and subcloned into the appropriate vectors. For pan-neuronal *C. elegans* expression, PiGM-Iq expression cassette and the corresponding control cassette were inserted into the JB6-Psnb1 vector (Gift from Dr. Zhitao Hu, Queensland Brain Institute). pCFJ90-Pmyo-2::mCherry::unc-54utr co-injection plasmid for *C. elegans* was also acquired from D. Zhitao Hu (Queensland Brain Institute). For LexA-inducible expression in *Drosophila, pJFRC18-8XLexAop2-RGS2(Δ1-53)::CRY2(PHR)::T2A::CIBN::eGFP::CaaX* and *pJFRC18-8XLexAop2-RGS2(Δ1-53)::CRY2(PHR)(D387A)::T2A::CIBN::eGFP::CaaX* were generated by cloning the PiGM cassette into pJFRC18-8XLexAop2-mCD8::GFP ^26^ cut with XhoI and XbaI (NEB). For zebrafish expression, PiGM-Iq and the control expression cassettes were subcloned into the ptol2PSA_UAS expression plasmid.

### HEK293 cell culture, transfection and transduction

HEK293A cells (Thermo Fisher Scientific, MA, USA) were maintained in DMEM with 1g/L of D-glucose supplemented with 8% Fetal Bovine Serum/FBS and 1% Pen/Strip (Thermo Fisher Scientific, MA, USA and MilliporeSigma, MA, USA). For imaging experiments, cells were plated on glass coverslips and transfected with X-tremeGENE 9 or polyethylenimine/PEI (MilliporeSigma, MA, USA). Cells were used for experiments 2 days post transfection. For generating stable O-GECO1 cells, HEK293A cells were co-transfected with the pLenti-CMV-O-GECO1, pMDG.2 and psPAX2 and the media of the transfected cells were transferred to untransfected HEK293A cells after 48 and 72 hours. The pMDG.2 and psPAX2 helper plasmids were gifts from Professor Didier Trono, Federal Institute of Technology, Lausanne.

### Calcium and localization imaging

HEK293 cells stably expressing O-GECO1 are plated on glass coverslips and placed in an imaging chamber of an upright Olympus BX51WI microscope (Olympus, Tokyo, Japan) equipped with Hamamatsu Orca Flash 4.0 camera (Hamamatsu Photonics, Shizuoka, Japan) and Excelitas X-cite 110LED (Excelitas Technologies, MA, USA) light source. Calcium imaging of O-GECO1 were done with a water immersion 20x/NA0.5 objective (Carl Zeiss AG, Oberkochen, Germany) and a TRITC filter set (FF01-543-22 excitation filter, 562-Di03 dichroic filter and FF01-593/40 emission, Semrock, VT, USA). EGFP was imaged with a GFP filter set (472/30 excitation filter, FF495-Di03 dichroic filter and FF01-520/35 emission filter, Semrock). For localization experiment, a water immersion 40x/NA0.8 objective was used (Olympus, Tokyo, Japan). Imaging was performed with Micromanager 1.4.22 software (https://micro-manager.org/). For blue light stimulation, a 470nm LED with 9° lens (Luxeonstar, Alberta, Canada) was placed under the condenser of the microscope and controlled by a LED controller via its associated LED Driver Control Panel V3.1.0 software (Mightex, Ontario, Canda). Most experiments were done with the stimulation of light intensity of of 0.22mW/mm^2^ and 1s duration. Cells were placed in HEPES-buffered extracellular solution (in mM: 140 NaCl, 3 KCl, 10 HEPES, 1 MgCl_2_, 2 CaCl_2_, 10 glucose, pH 7.4 and 290-310mOsm/L). Drugs was directly perfused into the imaging chamber.

### GloSensor cAMP assay

HEK293 cells were seeded in 96 well plate coated with poly-_L_-lysine and transfected with the test construct, pcDNA3-GloSensor (Promega, WI, USA) and pcDNA3-D_2_R-expression vector (generated with RT-PCR of rat brain mRNA library). Cells were incubated with 2mM luciferin (Cayman Chemicals, MI, USA) for 2 hours at room temperature prior to experiment. Quinpirole and isoprenaline (MilliporeSigma, MA, USA) were added to the well during the experiment to achieve the final concentration of 5 ρM and 10 respectively. Luminescence was recorded every minute (1s integration time) on a Tecan Spark microplate reader (Zurich, Switzerland).

### Image analysis and statistical analysis

The image stacks were analyzed with ImageJ 1.50. For O-GECO1 measurement, the cytosolic regions were traced manually and the mean intensity values for each cell were measured along with a non-fluorescent region of the field of view which is used as background. Background values were subtracted for each frame and the F_0_ value was determined as the mean of ten values immediately preceding blue light stimulation. For each frame, the F_0_ value was subtracted from the mean cytosolic intensity for each cell and the resulting value was normalized to the F_0_ value.

Statistical comparisons were performed with Graphpad Prism 8.3 using the indicated tests (Graphpad, CA, USA).

### *Caenorhabditis elegans* generation and mobility measurement

PiGM-Iq expression cassette was subcloned into a JB6-Psnb1 plasmid with pan-neuronal *Caenorhabditis elegans* synaptobrevin-1 promoter. The transgeneic animals were generated via microinjection into the gonad with a pCFJ90-pMyo-2::mCherry::unc-54utr co-injection marker and selected for appropriate crosses. Expression plasmids were maintained extra-chromosomally.

For mobility measurement, staged adult hemaphrodites were transferred to a pre-seeded NGM plates and allowed to habituate for 3 minutes. Once habituated, locomotion was recorded for 1 minute (QIClick Digital CCD Camera, Teledyne Photometrics, AZ, USA). The plate was transferred to a transilluminator (BlueView MBE-300, Major Sciences, Taoyuan City, Taiwan) and illuminated for 30 seconds (470 nm, 0.30 mW/mm^2^). Locomotion was then recorded 1 minute post-illumination for a total of 1 minute. Proceeding this, worms were incubated in the dark for 20 minutes at 20°C, after which another set of pre-and post-illumination locomotion measurements were performed as described above. Body bends per minute were calculated by counting the number of flexions that occurred anterior to the pharynx as the worm moved forward. Movement was not counted during spontaneous reversals, and the flexion occurring directly after this movement was ignored. Although the experimenter was not blinded during the recording of the videos, the experimenter was blinded for the analysis of the video for body bend quantification.

### *Drosophila melanogaster* lines and germline transformation

Fly strains used were wild-type Canton-S, the mushroom-body-specific *R13F02-LexA* driver line ^43^, and the pan-neuronal *R57C10-LexA* driver line ^26, 44^. The LexAop-PiGM-Iq and LexAop-PiGM[D387A]-Iq lines were generated through injection of y^1^sc^1^v^1^P{nos-phiC31\int.NLS}X; P{CaryP}attP40; + embryos as previously described ^45^.

### Courtship conditioning assay for *Drosophila melanogaster*

Courtship conditioning was performed as described previously ^29, 46^. Briefly, training and testing were performed at room temperature (21°C ± 2°C) with no humidity control. Test males and virgin flies for testing were isolated immediately following eclosion and aged separately for 5-7 days. Test males were housed in groups no larger than 4 per vial to limit male-male aggression. Test males were housed in vials partially protected from light to limit activation of the tool by the LEDs of the incubators during aging. For the generation of mated females, virgin females were housed with males for 5 days to ensure mating. Courtship repression training was then conducted with transgenic male flies within 1 hour of the initiation of the light-cycle to ensure maximal activity and consistency. Mated females were mildly anaesthetised using CO_2_ and placed into a 0.5ml Eppendorf tube containing a moist tissue to maintain humidity. Once the mated female had fully recovered, test males were gently aspirated into the tube containing a mated female (trained) or alone (naïve). Flies were then either illuminated with pulsed blue light or left in ambient room lighting for 1 hour. Blue light illumination was delivered via a custom LED array (470 nm) for 10 seconds every 5 minutes for the duration of the training period. Males assigned to the trainer group who did not attempt to court the mated female within 10 minutes from the initiation of training were excluded from further testing.

Following training, males were gently aspirated into the test chamber, consisting of a 24-well cell culture plate sealed with parafilm, and left to acclimatise for 10 minutes. A virgin female was then aspirated into the chamber containing the male. Addition of the female initiated the testing period, and the male was filmed for 10 minutes. These videos were then manually analysed for courtship behaviours by an individual blinded to the lighting condition and training group assigned to that male. The Courtship index (CI) represents the proportion of time spent courting during the test period. Trials where little to no courtship attempts were initiated by the naïve male were removed on account of environmental and housing issues. Learning index (LI) of male flies in each condition was then calculated using the following formula: LI = (CI_naïve_ - CI_trained_)/CI_naïve_, where CI_naïve_ represents an average CI score calculated from 12-24 naïve males tested against decapitated virgins for each genotype.

### Confocal microscopy of *Drosophila* brains

Adult brains (1 day post eclosure) were dissected in PBS and fixed in PBS + 0.3% TritonX-100 (PBST) with 4% (v/v) paraformaldehyde (ProSciTech) for 20 mins before three washes in PBST. Brains were stained with the mouse anti-Bruchpilot (Brp) antibody (nc82, Developmental Studies Hybridoma Bank) at 1/40 dilution in PBST overnight, washed 3x PBST, counter-stained with an anti-mouse alexa-594 secondary antibody (1/1000) overnight. Brains were washed a final three times in PBST, mounted in Vectashield + DAPI (Vector Labs), and imaged under an Olympus FV3000 confocal microscope at 20x magnification.

### DRG neurons and growth cone turning assay

DRG neuronal cultures were prepared and imaged as previously described ^31, 47^. In brief, thoracic and lumbar dorsal root ganglia neurons were extracted from E17.5-E18.5 rat embryos and plated on glass coverslip coated with poly-_L_-ornithine (Sigma-Aldrich, MO, USA) for 4-6 hours before experiments.

Imaging of the growth cone were done on an inverted phase-contrast microscope (Eclipse TiE; Nikon Instruments, NY, USA) with a 100× oil immersion (1.40 NA) objective at 26-32°C. Mcropipettes (opening diameter of 0.5-0.8 μm) were filled with solution containing serotonin positioned at a 45° angle and 85 μm distance from the growth cone and a pulsatile ejection (5 psi, 1Hz; Picospritzer, Parker, CO, USA) was used to apply serotonin treatment. Photostimulation was performed using a digital mirror device (Mightex, Ontario, Canada) with 470 nm LED light on the side of serotonin delivery (2 s pulse every 5 min, 10 % intensity). Images were acquired with Evolve EMCCD camera (Teledyne Photometrics, AZ, USA) for 30 min (7 s interval for 30 min) with custom software (MatLab, MA, USA). For the analysis, turning angle was defined as the change in axon trajectory of the distal 10 μm of axon using NIH ImageJ (MD, USA).

### Generation of transgenic *Danio rerio*

PiGM-Iq or the light insensitive D387A expression cassettes were subcloned into the ptol2P2A_UAS vectors resulted in ptol2P2A_UAS_PiGM-Iq and ptol2P2A_UAS_PiGM-Iq(D387A). In some cases, the cDNA for mCherry were inserted in-frame between the CRY2PHR and the 2A sequences. The ptol2 constructs (50 ng/μl) were co-injected with Tol2 transposase mRNA into one-cell stage eggs from Tg(Tph2_Gal4)y298Et fish. The resulting zebrafish lines that express PIGM-Iq and PIGM-IqD387A constructs in serotonergic neurons are labelled as Tg(Tph2_Gal4)y298Et; UAS_PIMG-Iq and Tg(Tph2_Gal4)y298Et; UAS_PIMG-Iq(D387A).

Zebrafish embryos were illuminated with an IO Rodeo midi blue LED transilluminator (Smart Lab Technology) for periods of 2 hr every 5 min for 10 s of continuous illumination. PiGM-Iq mutants and PIGM-Iq(D387A) light insensitive mutants were stimulated with LED blue light simultaneously. Light intensity was measured as 0.021 mW/mm^2^.

To assess coiling behaviour at 22-26 hpf, PiGM-Iq and PiGM_Iq(D387A) embryos were placed in a petri dish to acclimatise for 30 s. Spontaneous coiling behaviour was recorded at a rate of 40 fps (frames per second) for a period of 2 min. To assess startle-response at 36 hpf, 48 hpf, 72 hpf, PIGM-Iq and PIGM_Iq(D387A) embryos were stimulated using a fine micro-capillary tip applied to the tail. Touch-startle response was recorded at 89 fps. In both assays, PiGM-Iq or PiGM-Iq(D387A) embryos were alternated during each experiment. Coiling and startle behaviours were recorded with Mightex super speed CCD camera (Scitech) on a SZX19 dissection microscope (Olympus, Japan) and the software SS Classic Camera App. All videos were analysed in Fiji and manually counted. For anatomical analysis of the embryos, confocal stacks were acquired in 25 μm increments covering total zebrafish thickness, using Olympus Confocal FV3000 Upright equipped with a x20 water immersion objective (1.25 NA).

## Author contributions

JYL, JLL, LF and OM RG conceived the work. J. Lockyer and J.Y. Lin designed and generated the PiGM-I constructs. JLL, AR, SV, RG, LF and CD performed experiments. JLL, AR, SV, JYL, RG, LF, CD and OM analysed the data. JYL, SV, JLL and OJM wrote the manuscript.

## Acknowledgements

The project was supported by NIH BRAIN Initiative U01NS090590 grant and NHMRC APP1103034 project grant to J.Y. Lin and NHMRC project grants APP1128784 and APP185220 and Ian Potter Foundation grant 20190091 to O.J.M. J.Y. Lin was additionally supported by ARC Future Fellowship FT160100056. C. S. Vicenzi, R. Gasperini and L. Foa were supported by NHMRC 1165616. We thank John McMullen for technical assistance.

---

**Supplementary figure 1.**
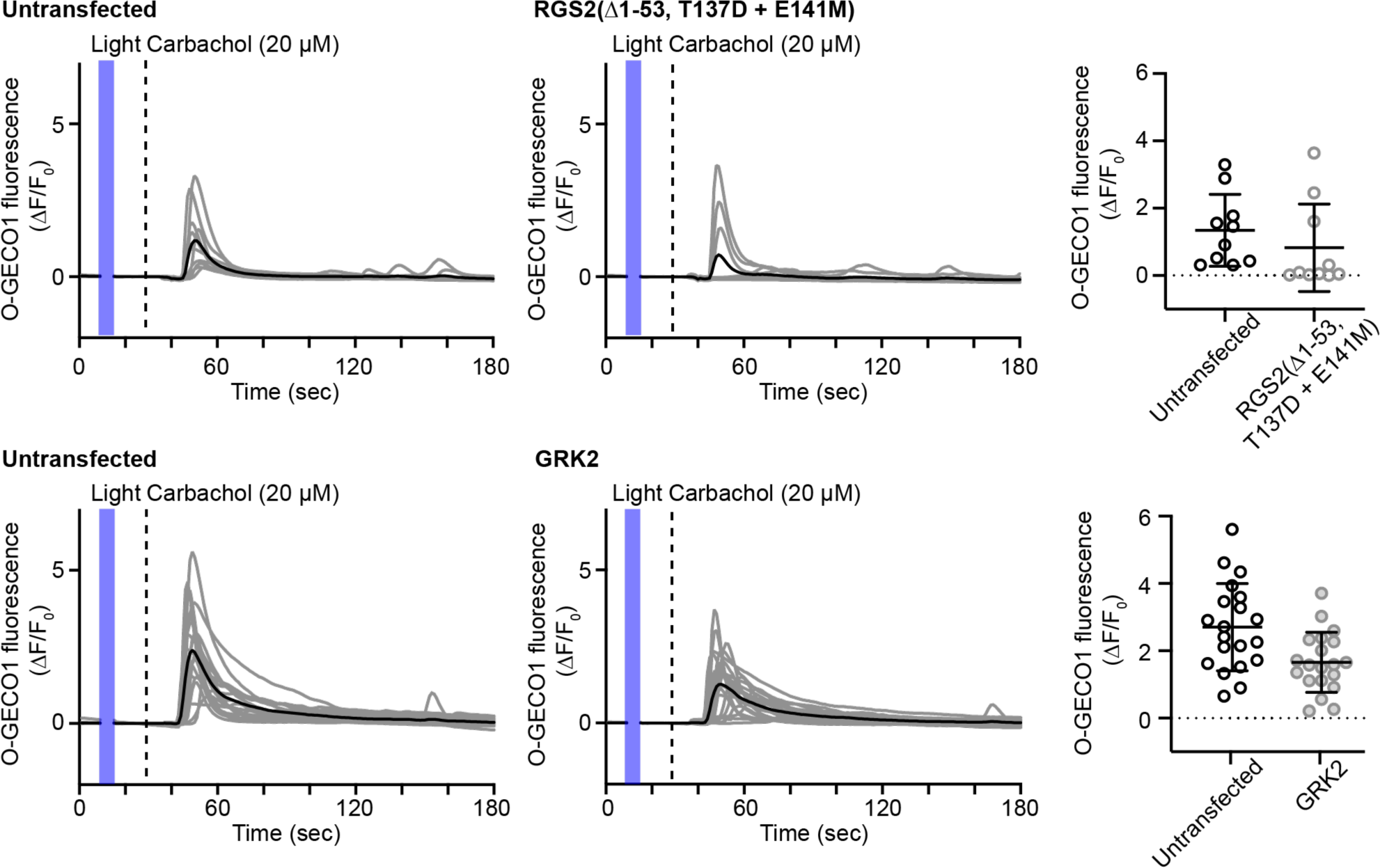
The carbachol-induced calcium elevation in cells expressing RGS2(Δ1-53)-CRY2PHR with T137D and E141M mutations and RGS domain of GRK2. The carbachol-induced calcium elevation in individual untransfected cells and cells expressing the respective constructs. Gray lines are the responses of individual cells and black lines are the mean responses. Although both constructs reduced the calcium response on average, the suppression is less efficient compared to PiGM-Iq.

**Supplementary figure 2.**
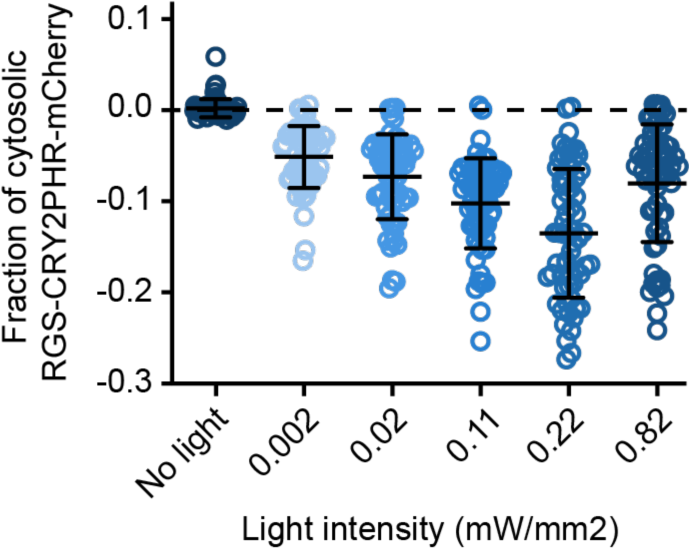
Cellular translocation RGS2(**Δ**1-53)-CRY2PHR-mCherry to the membrane CIBN at different light intensities. Fraction of cytosolic RGS2(Δ1-53)-CRY2PHR-mCherry when cells were stimulated with different light intensities. Maximum translocalization is observed with light intensity of 0.22 mW/mm^2^ whereas higher light intensity of 0.82 mW/mm^2^ reduces the fraction translocated to the membrane. All light intensities have significance level < 0.001 when compared with no light stimulation conditions. Significances were tested with Kuskal-Wallis test followed by Dunn’s multiple comparison tests against no light stimulation.

**Supplementary figure 3.**
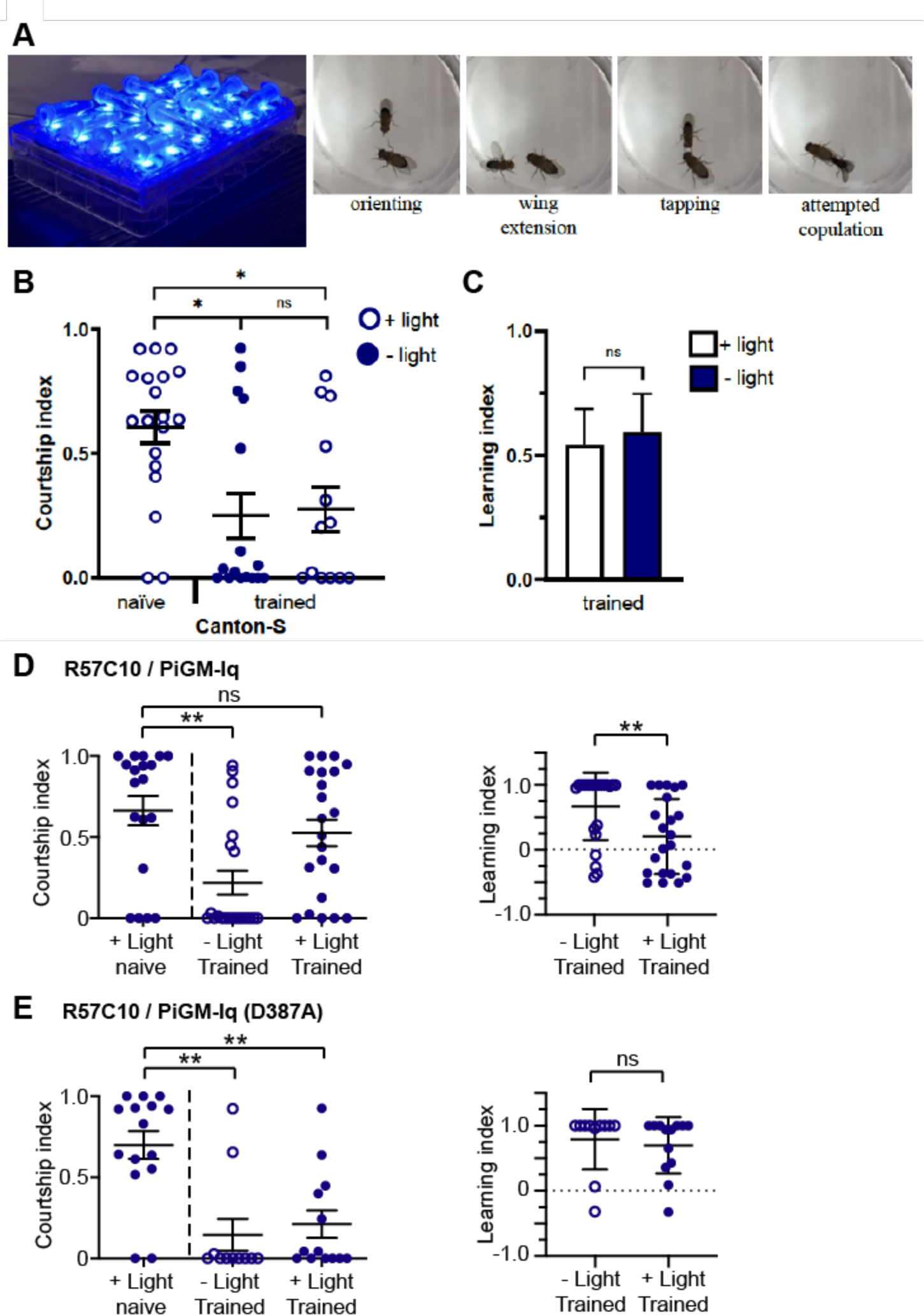
Testing the courtship conditioning assay in *D. melanogaster* and *D. melanogaster* with pan-neuronal PiGM-Iq expression. (A) 5-7 day old, sexually naïve males were trained with pre-mated females for 1 hour and pulsed with 470 nm light for 10 seconds every 5min. Males were then transferred to testing chambers containing a virgin female and their courtship behaviour was scored over a 10-minute period. (B) courtship index of male canton-S flies, Kruskal-Wallis test, ns indicates P >0.05, * indicates P < 0.05. (C) Learning index of male canton-S flies, Mann-Whitney test, ns indicates P >0.05. N = 13-19 flies/condition. (D) The courtship (left) and learning (right) indices of male flies expressing PiGM-Iq pan-neuronally (R57C10) showing the inhibition of learning. (E) Pan-neuronally expression of the light insensitive PiGM-Iq(D387A) mutant did not have light-induced disruption of learning. Error bars indicate SD.

**Supplementary material 1.**
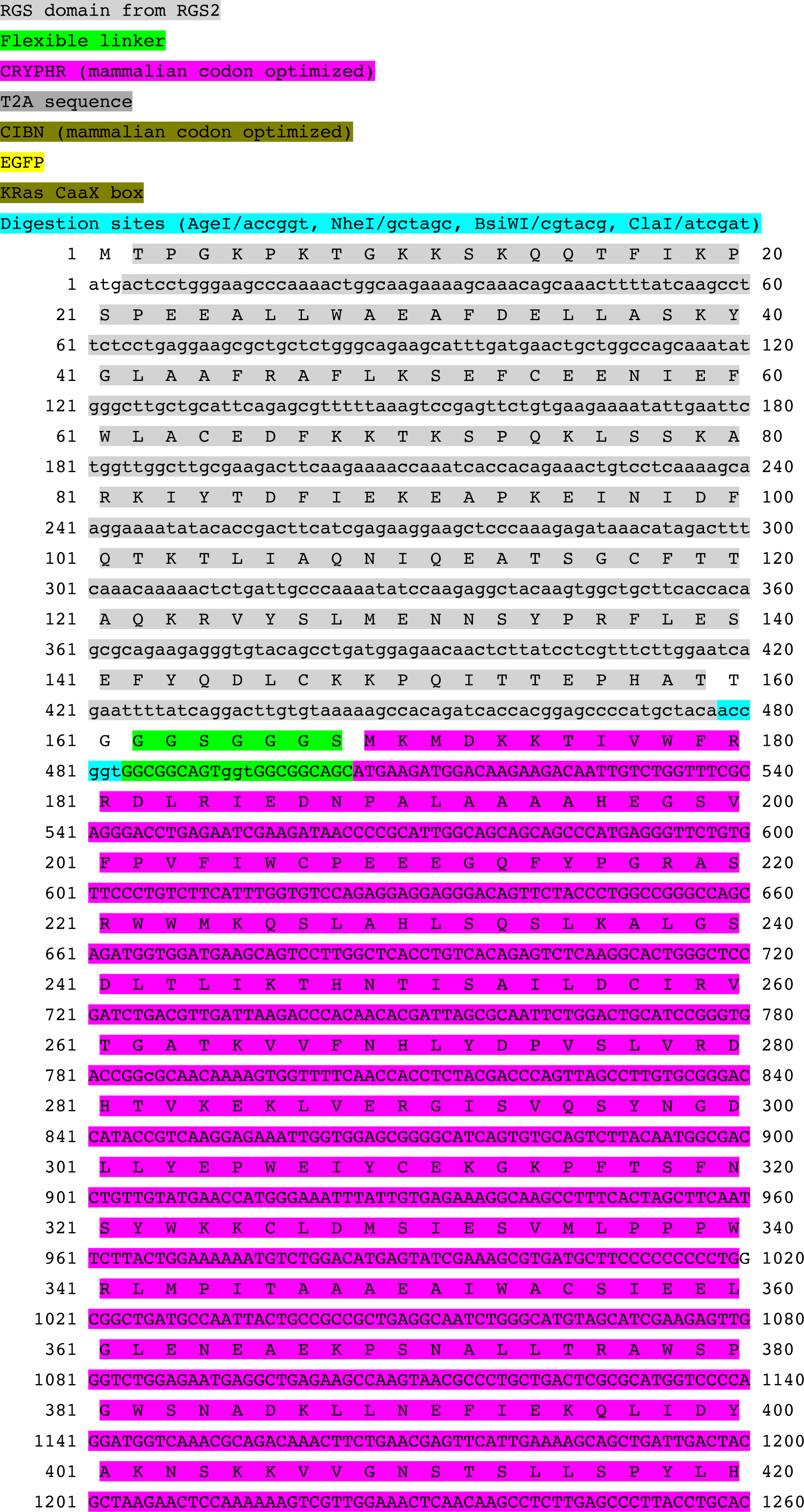
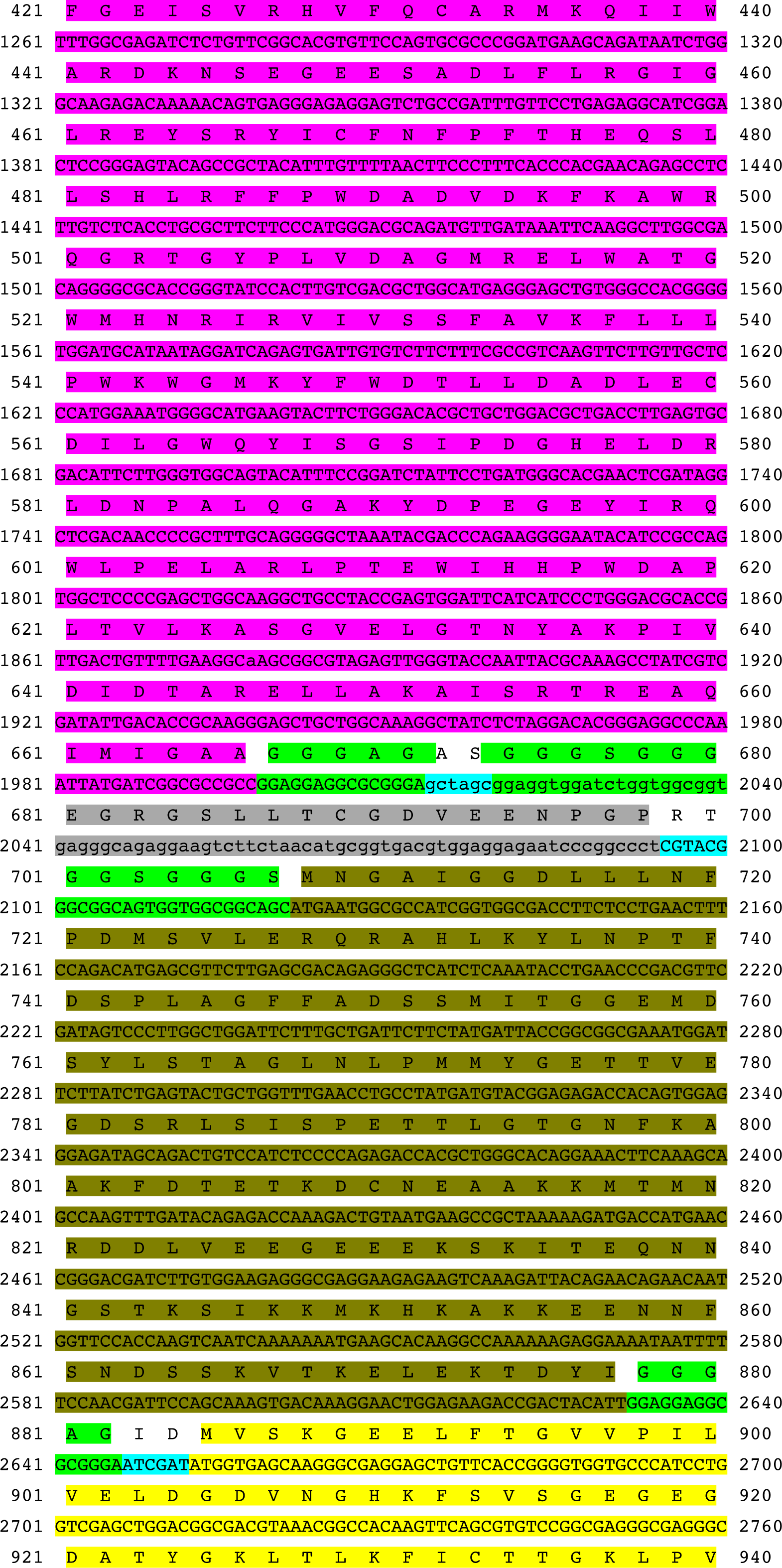
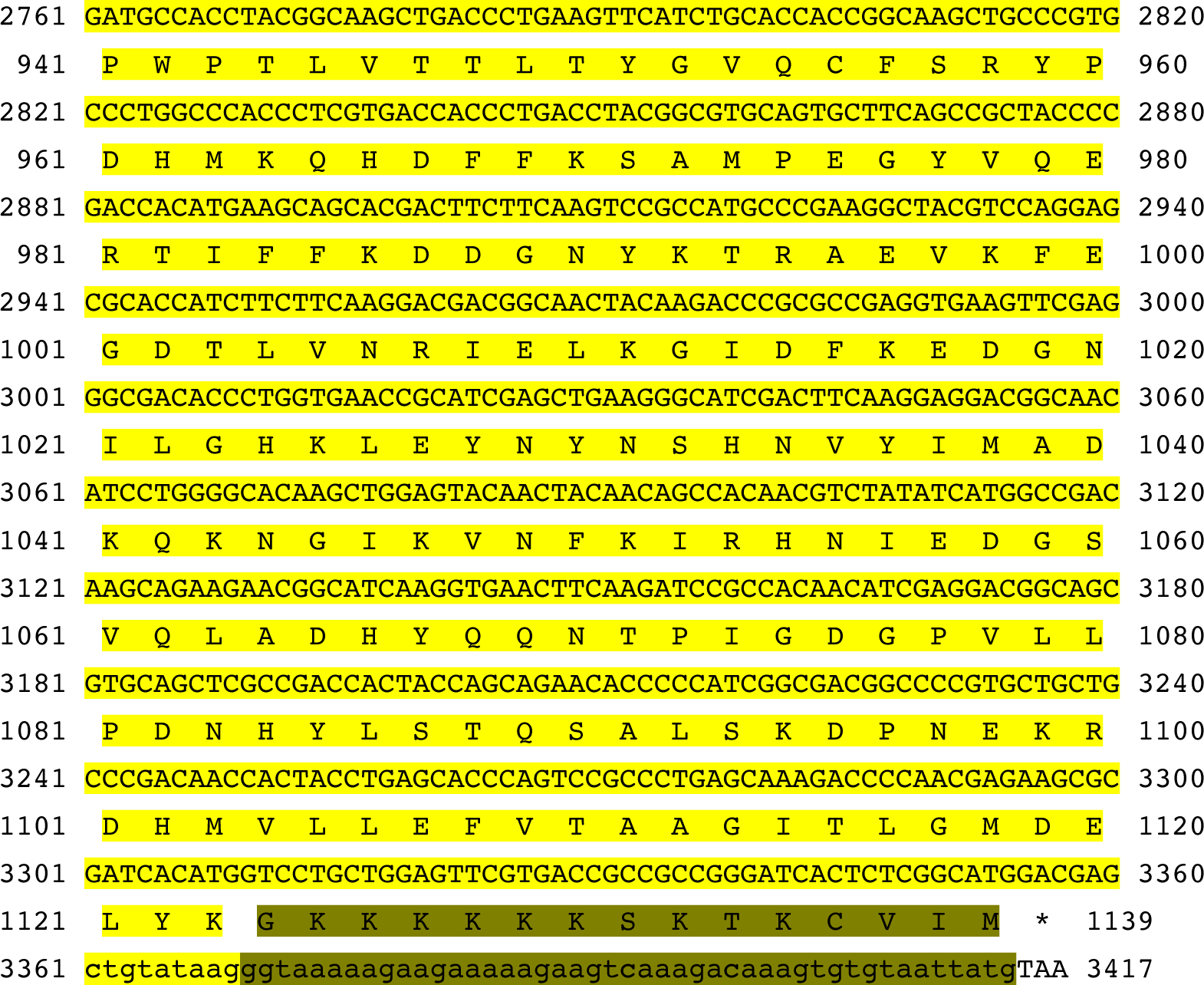
The annotated cDNA and amino acid sequences of final PiGM-Iq design.

**Supplementary material 2.**
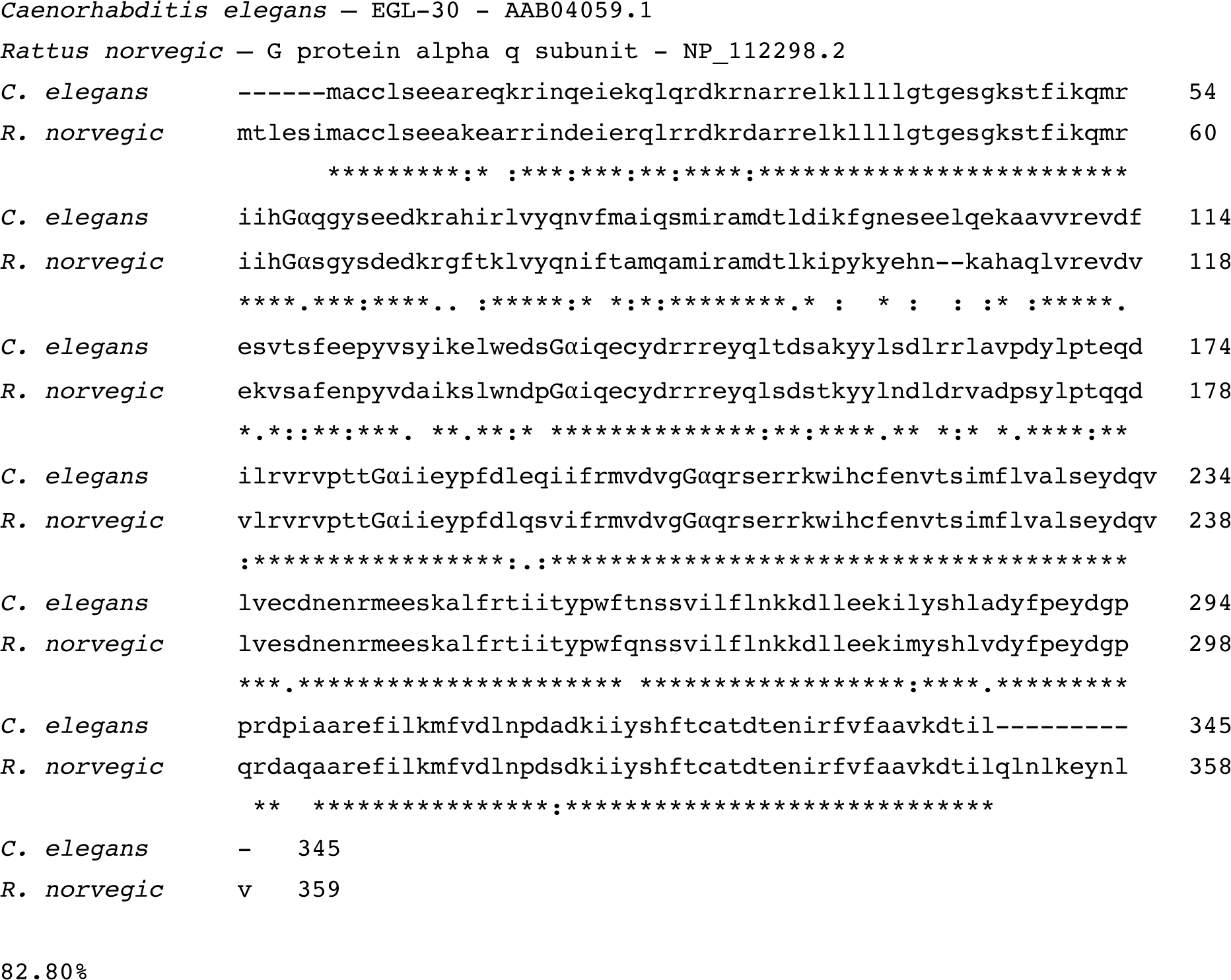
The amino acid sequence alignment and comparison of the *Caenorhabditis elegans* Gα_q_ homologue EGL-30 to the rat Gα_q_ used in this study.

**Supplementary material 3.**
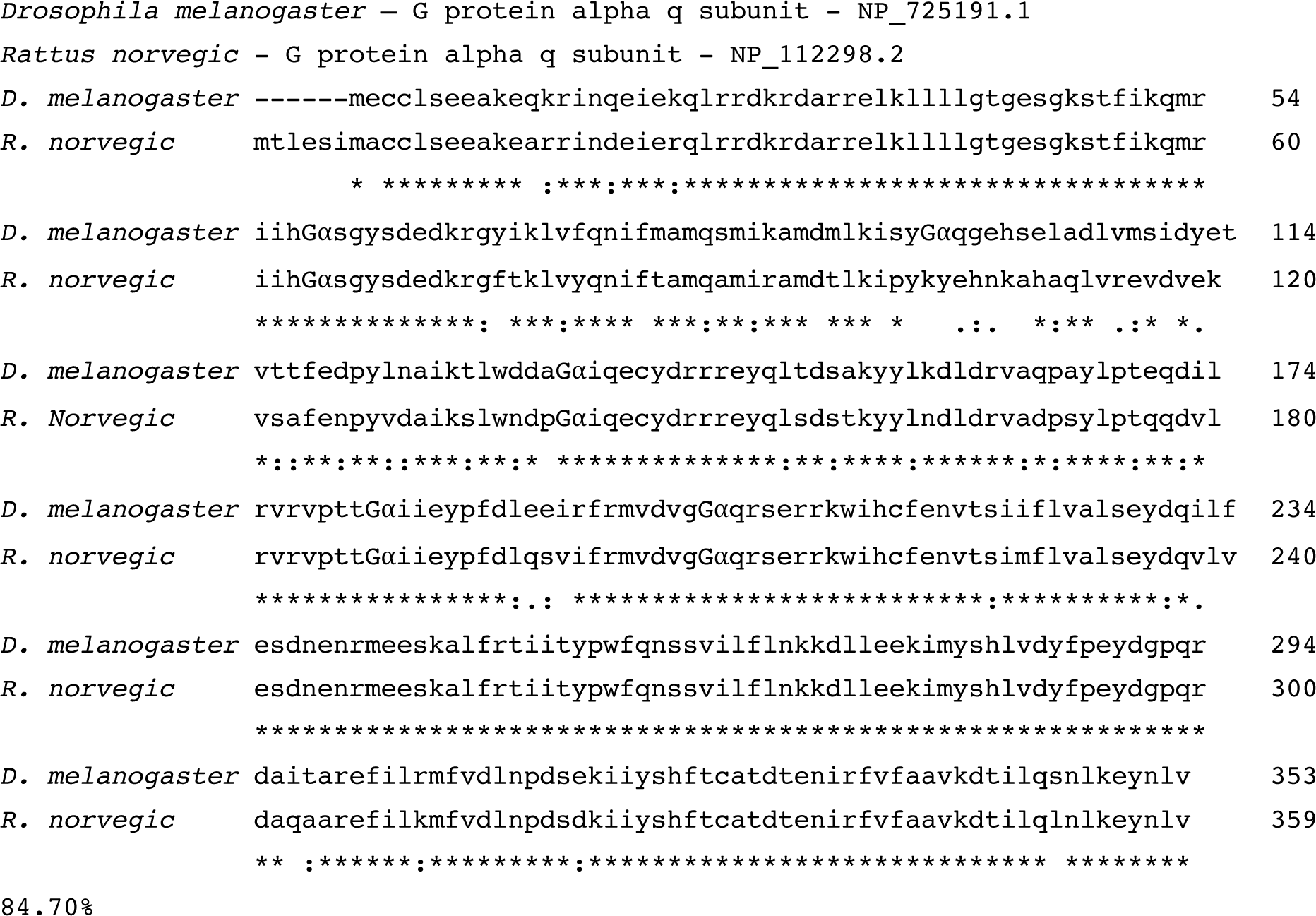
The amino acid sequence alignment and comparison of the *drosophila melanogaster* Gα_q_ homologue to the rat Gα_q_ used in this study.

**Supplementary material 4.**
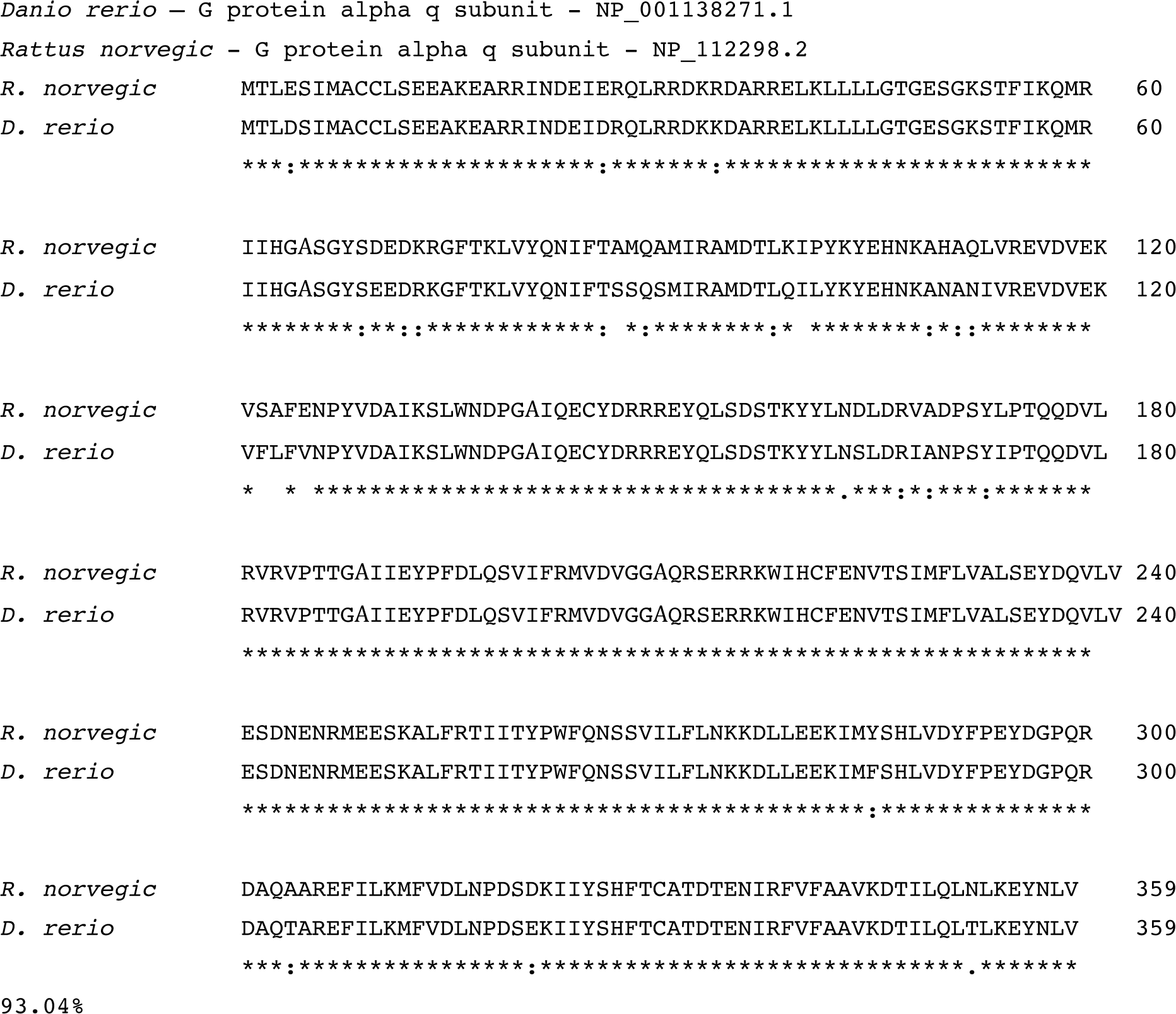
The amino acid sequence alignment and comparison of the *Dario rerio* Gα_q_ homologue to the rat Gα_q_ used in this study.

**Supplementary Table 1.** Quantification of *C. elegans* movements expressing PiGM-Iq before and after light stimulation.

|  | Movement (body-bends / minute) |  |  |
| --- | --- | --- | --- |
|  | N2 / wild-type (n = 12) | PiGM-lq (n = 11) | PiGM-lq(D387A) (n = 12) |
| Baseline | 40.25 ± 7.39 | 40.73 ± 9.66 | 33.33 ± 5.44 |
| After light pulse 1 | 38.00 ± 10.42 | 30.73 ± 8.59 | 34.50 ± 7.80 |
| Recovery | 31.75 ± 10.61 | 37.55 ± 10.95 | 28.33 ± 7.52 |
| After light pulse 2 | 30.67 ± 11.09 | 21.82 ± 10.78 | 25.50 ± 7.08 |

Presented as mean ± S.D.

**Supplementary Table 2.** Quantification of *drosophila* courtship index.

|  | Wild-type<br>(Canton-S) | R57C10 /<br>PiGM-lq | R57C10 /<br>PiGM-lq<br>(D387A) | R13F02 /<br>PiGM-lq | R13F02 /<br>PiGM-lq<br>(D387A) |
| --- | --- | --- | --- | --- | --- |
| Naïve +<br>Light | 0.61 ± 0.28<br>(n = 19) | 0.66 ± 0.40<br>(n = 19) | 0.70 ± 0.33<br>(n = 15) | 0.75 ± 0.27<br>(n = 16) | 0.49 ± 0.38<br>(n = 18) |
| Trained -<br>Light | 0.25 ± 0.36<br>(n = 16) | 0.22 ± 0.34<br>(n = 22) | 0.15 ± 0.32<br>(n = 11) | 0.29 ± 0.39<br>(n = 21) | 0.22 ± 0.28<br>(n = 16) |
| Trained +<br>Light | 0.28 ± 0.32<br>(n = 13) | 0.53 ± 0.38<br>(n = 22) | 0.22 ± 0.30<br>(n = 13) | 0.60 ± 0.41<br>(n = 17) | 0.17 ± 0.32<br>(n = 15) |

Presented as mean ± S.D.

**Supplementary Table 3.** Quantification of *drosophila* learning index.

|  | Wild-type<br>(Canton-S) | R57C10 /<br>PiGM-lq | R57C10 /<br>PiGM-lq<br>(D387A) | R13F02 /<br>PiGM-lq | R13F02 /<br>PiGM-lq<br>(D387A) |
| --- | --- | --- | --- | --- | --- |
| Trained -<br>Light | 0.59 ± 0.60<br>(n = 16) | 0.67 ± 0.52<br>(n = 22) | 0.79 ± 0.46<br>(n = 11) | 0.62 ± 0.53<br>(n = 21) | 0.55 ± 0.57<br>(n = 16) |
| Trained +<br>Light | 0.55 ± 0.53<br>(n = 13) | 0.21 ± 0.58<br>(n = 22) | 0.70 ± 0.43<br>(n = 13) | 0.20 ± 0.55<br>(n = 17) | 0.66 ± 0.64<br>(n = 15) |

Presented as mean ± S.D.

## References

1. Huber, K.M., Kayser, M.S. & Bear, M.F. Role for rapid dendritic protein synthesis in hippocampal mGluR-dependent long-term depression. Science 288, 1254–1257 (2000).

2. Stoop, R. Neuromodulation by oxytocin and vasopressin. Neuron 76, 142–159 (2012).

3. Heximer, S.P., et al. G protein selectivity is a determinant of RGS2 function. J Biol Chem 274, 34253–34259 (1999).

4. Asli, A., Sadiya, I., Avital-Shacham, M. & Kosloff, M. “Disruptor” residues in the regulator of G protein signaling (RGS) R12 subfamily attenuate the inactivation of Galpha subunits. Sci Signal 11 (2018).

5. Taylor, V.G., Bommarito, P.A. & Tesmer, J.J. Structure of the Regulator of G Protein Signaling 8 (RGS8)-Galphaq Complex: MOLECULAR BASIS FOR Galpha SELECTIVITY. J Biol Chem 291, 5138–5145 (2016).

6. Xie, G.X. & Palmer, P.P. How regulators of G protein signaling achieve selective regulation. J Mol Biol 366, 349–365 (2007).

7. Airan, R.D., Thompson, K.R., Fenno, L.E., Bernstein, H. & Deisseroth, K. Temporally precise in vivo control of intracellular signalling. Nature 458, 1025–1029 (2009).

8. Bailes, H.J., et al. Optogenetic interrogation reveals separable G-protein-dependent and -independent signalling linking G-protein-coupled receptors to the circadian oscillator. BMC Biol 15, 40 (2017).

9. Li, X., et al. Fast noninvasive activation and inhibition of neural and network activity by vertebrate rhodopsin and green algae channelrhodopsin. Proc Natl Acad Sci U S A 102, 17816–17821 (2005).

10. Yu, G., et al. Optical manipulation of the alpha subunits of heterotrimeric G proteins using photoswitchable dimerization systems. Sci Rep 6, 35777 (2016).

11. Gasser, C., et al. Engineering of a red-light-activated human cAMP/cGMP-specific phosphodiesterase. Proc Natl Acad Sci U S A 111, 8803–8808 (2014).

12. Hannanta-Anan, P. & Chow, B.Y. Optogenetic Inhibition of Galpha(q) Protein Signaling Reduces Calcium Oscillation Stochasticity. ACS Synth Biol 7, 1488–1495 (2018).

13. O’Neill, P.R. & Gautam, N. Subcellular optogenetic inhibition of G proteins generates signaling gradients and cell migration. Mol Biol Cell 25, 2305–2314 (2014).

14. Kennedy, M.J., et al. Rapid blue-light-mediated induction of protein interactions in living cells. Nat Methods 7, 973–975 (2010).

15. Atwood, B.K., Lopez, J., Wager-Miller, J., Mackie, K. & Straiker, A. Expression of G protein-coupled receptors and related proteins in HEK293, AtT20, BV2, and N18 cell lines as revealed by microarray analysis. BMC Genomics 12, 14 (2011).

16. Wu, J., et al. Improved orange and red Ca(2)+/-indicators and photophysical considerations for optogenetic applications. ACS Chem Neurosci 4, 963–972 (2013).

17. Roy, A.A., et al. RGS2 interacts with Gs and adenylyl cyclase in living cells. Cell Signal 18, 336–348 (2006).

18. Salim, S., Sinnarajah, S., Kehrl, J.H. & Dessauer, C.W. Identification of RGS2 and type V adenylyl cyclase interaction sites. J Biol Chem 278, 15842–15849 (2003).

19. Sinnarajah, S., et al. RGS2 regulates signal transduction in olfactory neurons by attenuating activation of adenylyl cyclase III. Nature 409, 1051–1055 (2001).

20. Gu, S., et al. Unique hydrophobic extension of the RGS2 amphipathic helix domain imparts increased plasma membrane binding and function relative to other RGS R4/B subfamily members. J Biol Chem 282, 33064–33075 (2007).

21. Taslimi, A., et al. Optimized second-generation CRY2-CIB dimerizers and photoactivatable Cre recombinase. Nat Chem Biol 12, 425–430 (2016).

22. Day, P.W., et al. Characterization of the GRK2 binding site of Galphaq. J Biol Chem 279, 53643–53652 (2004).

23. Nance, M.R., et al. Structural and functional analysis of the regulator of G protein signaling 2-galphaq complex. Structure 21, 438–448 (2013).

24. Brundage, L., et al. Mutations in a C. elegans Gqalpha gene disrupt movement, egg laying, and viability. Neuron 16, 999–1009 (1996).

25. Lackner, M.R., Nurrish, S.J. & Kaplan, J.M. Facilitation of synaptic transmission by EGL-30 Gqalpha and EGL-8 PLCbeta: DAG binding to UNC-13 is required to stimulate acetylcholine release. Neuron 24, 335–346 (1999).

26. Pfeiffer, B.D., et al. Refinement of tools for targeted gene expression in Drosophila. Genetics 186, 735–755 (2010).

27. Szuts, D. & Bienz, M. LexA chimeras reveal the function of Drosophila Fos as a context-dependent transcriptional activator. Proc Natl Acad Sci U S A 97, 5351–5356 (2000).

28. Han, K.A., Millar, N.S. & Davis, R.L. A novel octopamine receptor with preferential expression in Drosophila mushroom bodies. J Neurosci 18, 3650–3658 (1998).

29. Zhou, C., et al. Molecular genetic analysis of sexual rejection: roles of octopamine and its receptor OAMB in Drosophila courtship conditioning. J Neurosci 32, 14281–14287 (2012).

30. Inagaki, H.K., et al. Optogenetic control of Drosophila using a red-shifted channelrhodopsin reveals experience-dependent influences on courtship. Nat Methods 11, 325–332 (2014).

31. Vicenzi, S., Foa, L. & Gasperini, R.J. Serotonin functions as a bidirectional guidance molecule regulating growth cone motility. Cell Mol Life Sci 78, 2247–2262 (2021).

32. McLean, D.L. & Fetcho, J.R. Ontogeny and innervation patterns of dopaminergic, noradrenergic, and serotonergic neurons in larval zebrafish. J Comp Neurol 480, 38–56 (2004).

33. Yokogawa, T., Hannan, M.C. & Burgess, H.A. The dorsal raphe modulates sensory responsiveness during arousal in zebrafish. J Neurosci 32, 15205–15215 (2012).

34. Hunt, T.W., Fields, T.A., Casey, P.J. & Peralta, E.G. RGS10 is a selective activator of G alpha i GTPase activity. Nature 383, 175–177 (1996).

35. Miyamoto, T., et al. Rapid and orthogonal logic gating with a gibberellin-induced dimerization system. Nat Chem Biol 8, 465–470 (2012).

36. Wu, H.D., et al. Rational design and implementation of a chemically inducible heterotrimerization system. Nat Methods 17, 928–936 (2020).

37. Benedetti, L., et al. Optimized Vivid-derived Magnets photodimerizers for subcellular optogenetics in mammalian cells. Elife 9 (2020).

38. Guntas, G., et al. Engineering an improved light-induced dimer (iLID) for controlling the localization and activity of signaling proteins. Proc Natl Acad Sci U S A 112, 112–117 (2015).

39. Zheng, B., et al. RGS-PX1, a GAP for GalphaS and sorting nexin in vesicular trafficking. Science 294, 1939–1942 (2001).

40. Lin, J.Y., Lin, M.Z., Steinbach, P. & Tsien, R.Y. Characterization of engineered channelrhodopsin variants with improved properties and kinetics. Biophys J 96, 1803–1814 (2009).

41. English, J.G., et al. VEGAS as a Platform for Facile Directed Evolution in Mammalian Cells. Cell 178, 748–761 e717 (2019).

42. Stoeber, M., et al. A Genetically Encoded Biosensor Reveals Location Bias of Opioid Drug Action. Neuron 98, 963–976 e965 (2018).

43. Vogt, K., et al. Shared mushroom body circuits underlie visual and olfactory memories in Drosophila. Elife 3, e02395 (2014).

44. Henry, G.L., Davis, F.P., Picard, S. & Eddy, S.R. Cell type-specific genomics of Drosophila neurons. Nucleic Acids Res 40, 9691–9704 (2012).

45. Marshall, O.J. & Brand, A.H. Chromatin state changes during neural development revealed by in vivo cell-type specific profiling. Nat Commun 8, 2271 (2017).

46. van Swinderen, B. & Hall, J.C. Analysis of conditioned courtship in dusky-Andante rhythm mutants of Drosophila. Learn Mem 2, 49–61 (1995).

47. Pavez, M., et al. STIM1 Is Required for Remodeling of the Endoplasmic Reticulum and Microtubule Cytoskeleton in Steering Growth Cones. J Neurosci 39, 5095–5114 (2019).

48. Han, B., Bellemer, A. & Koelle, M.R. An evolutionarily conserved switch in response to GABA affects development and behavior of the locomotor circuit of Caenorhabditis elegans. Genetics 199, 1159–1172 (2015).

